# Phylum-wide propionate degradation and its potential connection to poly-γ-glutamate biosynthesis in *Candidatus* Cloacimonadota phylum

**DOI:** 10.1101/2025.11.11.687785

**Authors:** Magdalena Calusinska, Malte Herold, Dominika Klimek, Marie Bertucci, Sébastien Lemaigre, Sébastien Cambier, Simone Zorzan, Céline Leclercq, Jan Dolfing, Maria Westerholm, Bettina Müller, Leila Nasirzadeh, Anna Schnürer, Paul Wilmes, Philippe Delfosse, Xavier Goux

**Affiliations:** Environmental and Industrial Biotechnology, Luxembourg Institute of Science and Technology, Hautcharage L-4940, Luxembourg; The Faculty of Science, Technology and Medicine (FSTM), University of Luxembourg, Esch-sur-Alzette L-4365, Luxembourg; Faculty of Energy and Environment, Northumbria University, Newcastle-upon-Tyne NE1 8QH, United Kingdom; Department of Molecular Sciences, Swedish University of Agricultural Sciences, BioCentre, Uppsala SE-75007, Sweden; Department of Biomedical and Clinical Sciences, The Division of Cell and Neurobiology, Linkoping University, Linkoping 581 83, Sweden; Luxembourg Centre for Systems Biomedicine, University of Luxembourg, Esch-sur-Alzette, L-4365 Luxembourg; Rectorate, Université du Luxembourg, Maison du Savoir, Esch-sur-Alzette L-4365, Luxembourg

**Keywords:** Anaerobic digestion, genomics, microbial isolation, syntrophic propionate oxidation

## Abstract

The candidate phylum Cloacimonadota is frequently detected in anaerobic environments such as anaerobic digestion (AD) reactors, hydrothermal vents, and deep-sea sediments, yet its metabolism remains poorly understood due to the lack of cultured representatives. Metagenomic evidence suggests capacities for amino acid fermentation, cellulose degradation, and production of carbohydrate-active enzymes, with particular interest in their presumed role in syntrophic propionate oxidation (SPO), a key bottleneck in AD. However, a complete methylmalonyl-CoA (mmc) pathway, central to SPO, has not been previously identified in Cloacimonadota genomes. Here, we report results from a lab-scale anaerobic baffled reactor fed with sugar beet pulp, where a sharp increase in an uncultured Cloacimonadota OTU coincided with recovery of methanogenesis and enhanced methane production. Metagenomic and metatranscriptomic analyses enabled metabolic reconstruction of this OTU, complemented by a curated database of 47 genome-resolved Cloacimonadota species. Comparative genomics revealed conserved protein clusters indicative of an alternative mmc pathway, suggesting that this variant of the SPO pathway is a widespread, phylum-specific trait potentially linked to protein degradation and poly-γ-glutamate biosynthesis. Network analysis identified the methanogenic archaeon Methanothrix as a primary syntrophic partner, an interaction further supported by propionate-fed enrichment cultures showing co-occurrence of Cloacimonadota and Methanothrix species. Our study sheds light on the Cloacimonadota metabolism, advancing our understanding of their ecological roles and potential for biotechnological applications.

## Introduction

The candidate phylum Cloacimonadota, formerly *Candidatus* Cloacimonetes, is a part of the *Fibrobacteres-Chlorobi-Bacteroidetes* superphylum. Cloacimonadota comprises gram-negative bacteria that are metabolically poorly characterized. Despite several isolation attempts, no pure cultures have been obtained, although successful enrichments have been reported (Dyksma et al., 2020; Pelletier et al., 2008). Cloacimonadota are predominantly found in anaerobic environments, often associated with methanogenic habitats such as anaerobic digestion (AD) reactors, deep-sea sediments, and hydrothermal vents (Calusinska et al., 2018; Johnson & Hug, 2022; Villanueva et al., 2021). In full-scale AD units, they typically constitute between 1% and 12% of the bacterial community (Calusinska et al., 2018). In deep-sea anoxic waters, Cloacimonadota accounts for between 5% to 15% of bacterial 16S rRNA gene reads (Villanueva et al., 2021).

The first genome sequence of *Ca*. Cloacimonas acidaminovorans was reconstructed over 15 years ago from a metagenomic library of a municipal wastewater treatment plant (WWTP), where it represented 2% of a fosmid library (Pelletier et al., 2008). Since then, multiple draft genomes of Cloacimonadota have been recovered from lab-scale and full-scale AD reactors (e.g. Broeksema et al., 2017; Dyksma et al., 2020; Dyksma & Gallert, 2019), hydrothermal vents (Villanueva et al., 2021), animal gut-associated habitats (Hervé et al., 2020; Marynowska et al., 2023), among others (Johnson & Hug, 2022). Based on the encoded metabolic potential, Cloacimonadota are recognized as amino acid-fermenting bacteria (Dyksma et al., 2020; Pelletier et al., 2008). Their ability to degrade cellulose has also been documented (Limam et al., 2014), and they possess a diverse repertoire of carbohydrate-active enzymes (CAZymes) (Johnson & Hug, 2022), establishing them as key hydrolytic bacteria within the AD community. However, the main interest in this uncultured bacterial phylum stems from its presumed role in syntrophic propionate oxidation (SPO), a key process that helps overcome one of the major bottlenecks to efficient methane production in AD reactors (Westerholm et al., 2022). Several studies have linked increased abundance of different Cloacimonadota species to propionate consumption (e.g., Dyksma & Gallert, 2019; Nobu et al., 2015), suggesting that SPO might be a common feature of the phylum. Conversely, decreases in their relative abundance have also been reported under conditions of elevated propionate levels (Singh et al., 2021). No reconstructed Cloacimonadota genome contains a complete set of genes encoding the key enzymes of the methylmalonyl-CoA (mmc) pathway, which is widely recognized as being involved in SPO. And the lack of cultured representatives from the Cloacimonadota phylum further complicates the elucidation of their role in the SPO process and currently limits the group to *Candidatus* status only.

This study was instigated by a lab-scale AD experiment in which a single operational taxonomic unit (OTU), affiliated with an uncultured Cloacimonadota organism, emerged in an acidified anaerobic baffled reactor (ABR). This emergence coincided with the spontaneous reestablishment of methanogenic conditions, likely driven by enhanced degradation of accumulated volatile fatty acids, which in turn enabled increased methane production at high organic loading rates (OLR) with sugar beet pulp (SBP) as feedstock (Fig. 1). To further characterize the enriched Cloacimonadota species, we sequenced the community metagenome and metatranscriptome, which allowed reconstruction of a complete genome for the Cloacimonadota OTU. We also compiled a database of Cloacimonadota genomes from both public repositories and our own datasets. Whole-genome phylogenetic analysis resolved 47 distinct genome-based species (GS) within the phylum. To clarify their role in AD, we compared the Cloacimonadota pangenome with genomes of other AD-associated microbes. Focusing on their potential function as propionate oxidizers, our analysis of conserved protein clusters (PCs) supports the presence of an alternative mmc pathway in Cloacimonadota. The widespread occurrence of this pathway indicates that propionate oxidation may represent a phylum-wide trait, potentially linked to protein degradation and poly-γ-glutamate (PGA) production. Correlation network analysis (CNA) further suggested that methanogenic archaea of the genus *Methanothrix* are the primary syntrophic partners of our Cloacimonadota OTU. Notably, this archaeon was also found co-enriched with another Cloacimonadota species in an enrichment culture aimed at isolating the first cultivated strain of the phylum, although successful isolation was not achieved.

**Fig. 1.**
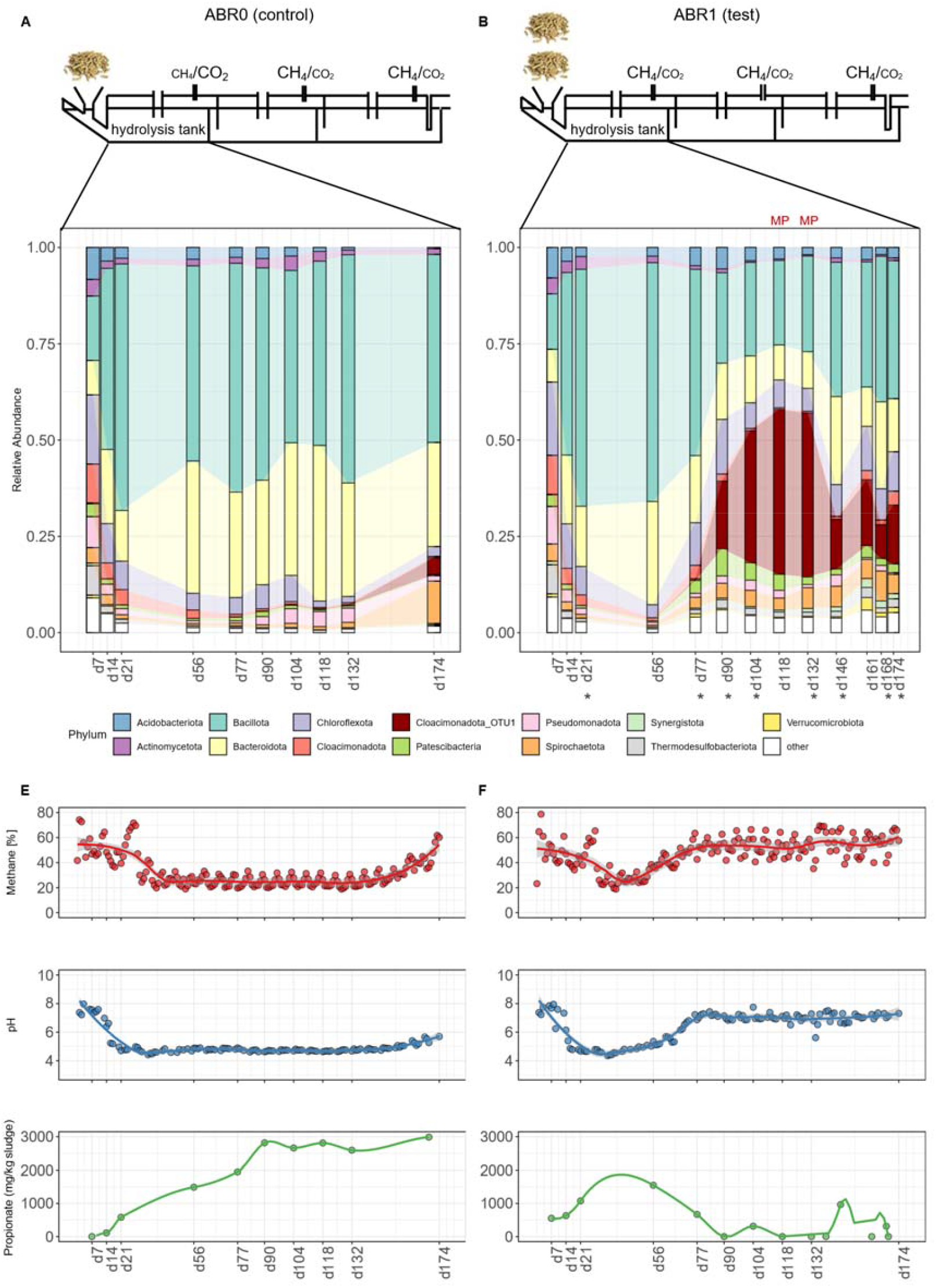
Overview of the Anaerobic Baffled Reactor (ABR) design and operation (A and B), the microbial communities established in the hydrolysis tank (C and D), and the recorded parameters (E and F) for the control and test ABR reactors (left and right panel, respectively). The abundance of Cloacimonadota OTU_1 is highlighted in the graphs. Samples analyzed for metagenomics (MG) and metatranscriptomics (MT) are marked with an asterisk and metaproteomics are indicated with MP.

## Material and methods

### Anaerobic baffled reactors: operation, sampling, and 16S rRNA gene amplicon sequencing

Two three-compartment anaerobic baffled reactors (ABR) were operated under mesophilic conditions for 174 days. Both reactors were inoculated with anaerobic sludge sourced from a full-scale methanogenic digester treating activated wastewater sludge, and samples were regularly collected from each ABR compartment. Further details are elaborated in the Supplementary Material (Methods 1.1). DNA and RNA macromolecules were co-extracted on the days indicated in Fig. 1, using the AllPrep DNA/RNA Mini Kit (Qiagen, Hilden, Germany) and following the manufacturer’s protocol. Bacterial and archaeal 16S rRNA gene amplicon libraries were prepared using a previously optimized Illumina-compatible sequencing approach (Calusinska et al., 2018). Correlation network analysis (CNA) was conducted on normalized 16S rRNA reads following the method initially developed by (De Vrieze et al., 2018) with further modifications. The complete protocol is detailed in Supplementary Material (Methods 1.2), and additional results are available in Supplementary Dataset 1 (Tables S1-S4).

### Metagenomics, metatranscriptomics and metaproteomics

Eight time points from the hydrolysis tank of ABR1 were selected for metagenomic sequencing (Fig. 1D). Sequencing libraries were prepared with the Nextera XT kit (Illumina) and sequenced using the Illumina NextSeq 500 platform (University of Luxembourg), generating 49.9 Gb of 150 bp metagenomic paired-reads. Omics samples were analyzed at time points corresponding to the occurrence of OTU_1 at high relative abundance. One sample (day 132) was additionally sequenced using long-read Nanopore approach (BaseClear, The Netherlands) producing 11 Gb. Short Illumina reads were quality trimmed in CLC Genomics Workbench vV23.0.5 Qiagen), using a phred quality score of 20, minimum length of 50 and allowing no ambiguous nucleotides. Quality-trimmed reads were assembled using the CLC’s de novo assembly algorithm in a mapping mode, using automatic bubble size and word size, minimum contig length of 1000, mismatch cost of 2, insertion cost of 3, deletion cost of 3, length fraction of 0.9, and similarity fraction of 0.95. Metatranscriptomics was conducted on the same eight samples as metagenomics (Fig. 1D), generating 82.1 Gb of 150 bp paired-reads. Library preparations and data analysis followed previously optimized in-house protocols (Calusinska et al., 2020). Canu-1.8 was used to process Nanopore reads and generate the assembly (Koren et al., 2017). Illumina metagenomic reads were mapped back to the Nanopore assembly, and binning and bin refinement, including re-assembly, were performed using MetaWRAP v1.3 (Uritskiy et al., 2018) which combined binning results of CONCOCT (Alneberg et al., 2014), MaxBin2 (Wu et al., 2016), and Metabat2 (Kang et al., 2019). 52 refined MAGs were retained with greater than 50% checkm (Parks et al., 2015) completeness and below 10% contamination (Supplementary Dataset 2, Table S5). Bin.36, corresponding to Cloacimonadota OTU_1, was re-assembled by extracting mapping Illumina and Nanopore reads as input for UniCycler (Wick et al., 2017), which resulted in a circular genome of 2.3 Mb. Functional and taxonomic annotation was performed for the reconstructed MAGs (see below). llumina MG and MT (after rRNA removal) reads were mapped back to the final MAG set with bwa mem v0.7.12 (Li & Durbin, 2009) and Nanopore reads were mapped with minimap2 v2.17-r941 (Li, 2018). Coverage and TPM values on MAG level were determined with coverM v0.7 (Aroney et al., 2025). For MT analysis, salmon v1.4 (Patro et al., 2017) was used to quantify the transcriptional abundance of predicted genes (including genes from MAGs and unbinned contigs) including sequence bias correction and the –meta option. Metaproteomics was applied to duplicate samples corresponding to the time point days 118 and 132 (Fig. 1) to confirm the presence of enzymes from the mmc pathway of Claocimonadota origin. Further details on the methods used in this study are described in Supplementary Material (Methods 1.3). Results corresponding to these methods are available in Supplementary Datasets 2, 3 and 4.

### Collection of Cloacimonadota genomes and phylogenetic analysis

The Cloacimonadota genome database was constructed by integrating Cloacimonadota-annotated genomes downloaded from NCBI in May 2020, genomes reconstructed in this study, and others (Brandt et al., 2020; Supplementary Dataset 2, Table S7). Additional, taxonomy assignment was performed using GTDB-Tk v2.4 and database release 220 (Chaumeil et al., 2020), resulting in 121 retained genomes, meeting the criteria for medium to high quality MAGs, e.g. completeness higher than 50% and contamination lower than 10%. All genomes underwent dereplication using dRep (Olm et al., 2017) with ‘gANI -sa 0.965 -nc 0.6’. Genome dereplication at the species level resulted in the formation of 47 genome clusters, hereafter referred to as genome-based species (GS; Fig. 2). Following the microbial species delineation criteria proposed by (Barco et al., 2020), we considered two nearly complete MAGs with an average ANI below 96.5% over at least 60% of the aligned genome to represent distinct microbial species. Once the representative strains are cultured, GS will likely form a novel species. A whole-genome phylogenetic tree was constructed using Phylophlan3 (Asnicar et al., 2020) and visualized with iTOL (Letunic et al., 2021). Average Nucleotide Identity (ANI) and Average Aminoacid Identity (AAI) were calculated for enriched GS using OrthoANI (Lee et al., 2016) and the enveomics suite (Rodriguez & Konstantinidis, 2016) respectively.

**Fig. 2.**
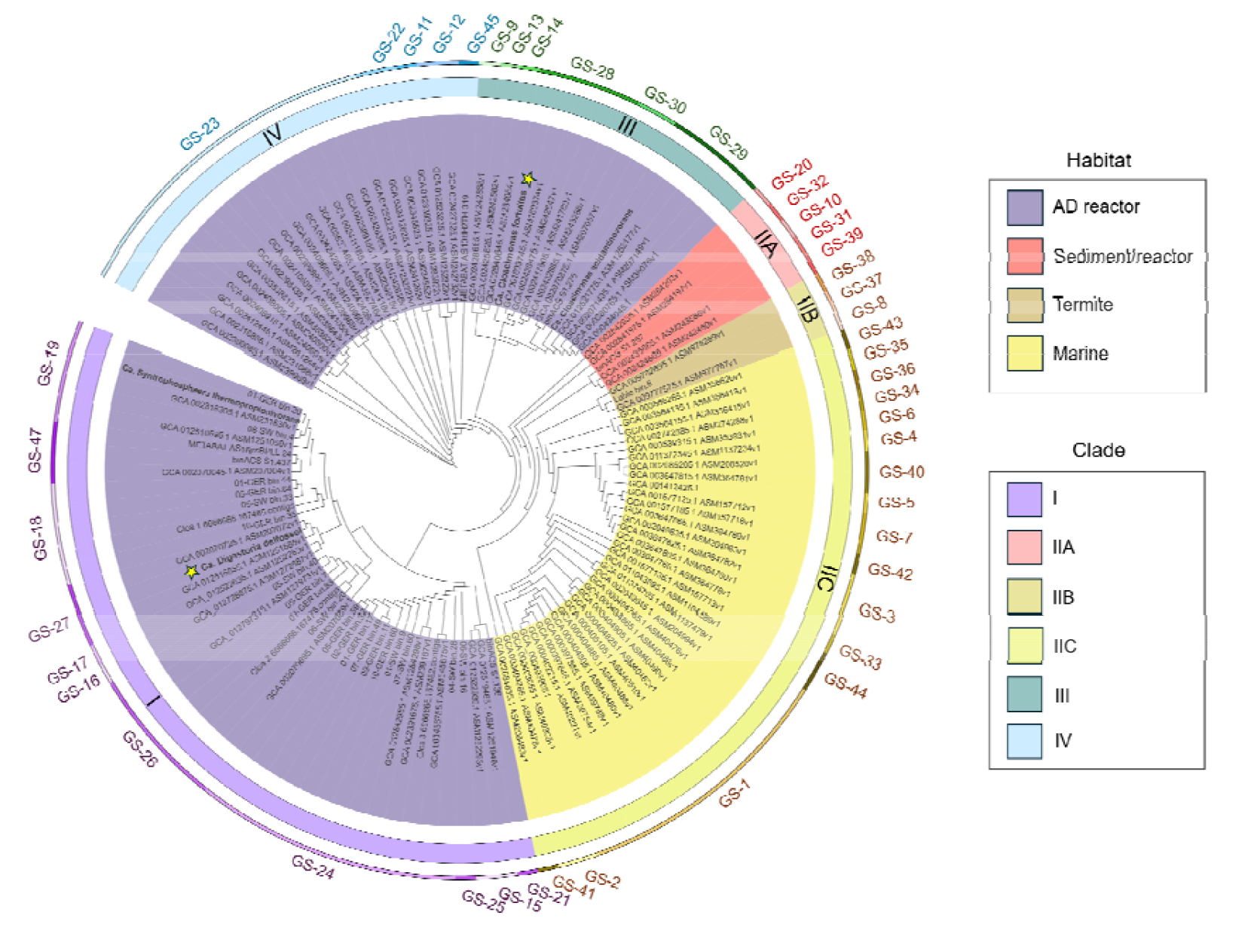
Phylogenetic placement of Cloacimonadota genomes and the resulting genome-based species (GS) based on a maximum likelihood phylogeny of concatenated conserved protein alignments. The phylogenetic positions of two candidate species identified in this study are marked with asterisks.

### Gene calling, functional annotation, and protein cluster generation

Gene calling was executed using Prodigal (Hyatt et al., 2010) within Prokka v1.14.6 (Seemann, 2014). All predicted proteins underwent functional annotation, including ortholog searches with emapper-2.0.1 and utilization of the eggNOG 5.0 database (Huerta-Cepas et al., 2019). Carbohydrate-active enzyme (CAZy) coding genes (CAZymes) were detected using dbCAN2 (Zhang et al., 2018) and the CAZy database v6 (Drula et al., 2022). Proteins were grouped into protein clusters (PCs) using MMseqs2 (Mirdita et al., 2019) with the parameters: ‘--min-seq-id 0.3 -c 0.8 --cov-mode 1’. Protein clusters were functionally annotated as described above. The functional assignment to the Kyoto Encyclopedia of Genes and Genomes (KEGG) orthologous categories (KOs) was done with GhostKoala (Kanehisa et al., 2016). Protein sequences were additionally annotated for Pfam domains using the HMMsearch against the Pfam HMM database (Mistry et al., 2021). Thermodynamics calculations of enzymatic reactions were made after (Dolfing, 2015). Gibbs free energy of formation values was based on (Hanselmann, 1991; Stephanopoulos et al., 1998; Thauer et al., 2008).

### Comparative analysis of the metabolic potential across major anaerobic digestion phyla

To characterize the metabolic potential of Cloacimonadota versus other major phyla present in the AD reactors, a previously generated database of MAGs was used (Campanaro et al., 2020). The KEGG and Pfam database were used to assign gene functionalities to the MAGs. To illustrate the specific functional roles of Cloacimonadota within the AD community, a non-metric multidimensional scaling (NMDS) projection was generated using Jaccard dissimilarities calculated from the Pfam matrix. This approach highlighted functional differentiation and potential metabolic contributions of Cloacimonadota in the AD ecosystem. The complete set of results is provided in Supplementary Dataset 4.

### Cloacimonadota isolation trials and genome reconstruction of enriched species

Due to the time lag between sample collection, data acquisition, and analysis, the isolate trials were conducted later than the ADR experiment, which had already been terminated. Therefore, we used the same seeding sludge as the ADR inoculum for the isolation experiment, rather than the reactor-enriched sludge. Given that poly-γ-glutamate (PGA) is a known virulence factor in other species and can confer nonspecific antibiotic resistance, sludge inoculum was incubated with broad-spectrum antibiotics (ampicillin, streptomycin, vancomycin, and ciprofloxacin) at the time of media inoculation, using modified DSM 621 and other custom media with various carbon sources. Enrichments were maintained under anaerobic conditions, in closed serum bottles for three months at either 37°C or room temperature with intermittent mixing. Further isolation efforts were designed to specifically target enriched Cloacimonadota. For a detailed description of the enrichment study design, see Supplementary Material (Methods 1.4) and Supplementary Dataset 5. To monitor the enrichment of Cloacimonadota, the enrichment cultures were periodically analyzed using Nanopore-based 16S rRNA gene sequencing. These 16S rRNA gene data are not included in the manuscript, as they were used solely to monitor the presence or absence of target taxa. One culture, initially amended with D-glucose, significantly enriched in Cloacimonadota and was subsequently passaged on modified DSM 621 medium supplemented with propionate (10 g.L^-1^) as the sole carbon source and incubated at room temperature with gentle mixing (80 rpm).

To enable genome reconstruction and further characterization of the species enriched in the examined propionate-fed culture, sequencing was performed on the Illumina MiSeq platform using the v3 reagent kit (2 × 250 bp paired-end reads), with library preparation carried out using the Illumina DNA Prep kit. The raw reads were quality-checked using FastQC and trimmed to remove adapters or low-quality bases and assembled into contigs using CLC Genomic Workbench. Genome binning was performed using MetaBAT2. CheckM was used to assess bin quality and GTDB-Tk was employed for taxonomic classification.

### Fluorescence *in situ* hybridization and microscopy

Fluorescence in situ hybridization (FISH) was employed to assess the laboratory culture enriched on propionate (10 g.L^-1^), initially confirming a significant enrichment of Cloacimonadota as indicated by 16S rRNA gene amplicon sequencing and metagenomics. To specifically visualize the presence of Cloacimonadota in this culture, we designed probes CloaPFishV1 (5’ GC<u>A</u>CCATGATGTTGCG 3’) and CloaOFishV2 (5’ GCG<u>C</u>CATGATGTTGCG 3’) which were utilized in equimolar ratio. These probes, differing by a single nucleotide, were specifically tailored to target any Cloacimonadota species within the tree clades I, III and IV identified in our study (Fig. 2). The design of these probes relied on aligned partial 16S rRNA gene sequences of Cloacimonadota from the study by (Calusinska et al., 2018), and their validation was conducted using the FISH probe search tool implemented in SILVA (https://www.arb-silva.de/search/testprobe/). The protocol is outlined in Supplementary Material (Methods 1.5). In addition, DAPI staining was used to label bacterial DNA and to visualize the PGA surrounding the Cloacimonadota flocs, following the method described by Szczepanek et al., 2002, which demonstrated that DAPI can be used for in situ detection of PGA. Gram-staining protocol was applied to the PBS-washed culture and observed under light microscopy (Zeiss).

## Results and discussion

### Novel propionate-consuming Cloacimonadota OTU_1 enriched in reactors operated with high organic loading rates

In this study, we operated two tri-compartment ABR reactors to evaluate the effect of physical separation of AD stages on methane production. ABR1 (test reactor) was fed with gradually increasing OLR of SBP, while ABR0 (control reactor) received cautiously regulated feed. Due to highly fermentable nature of the substrate, the 1^st^ compartment, designated as the hydrolysis tank became acidified early in the experiment for both reactors, regardless of the applied OLR. Notably, after day 56, a sharp increase in the abundance of a distinct Cloacimonadota species (OTU_1) was observed in the hydrolysis tank of ABR1, coinciding with a rapid onset of propionate degradation (Fig. 1; Supplementary Dataset 1, Table S1 – S3). This led to a gradual decline in propionate concentration from 1550 mg.kg_sludge_ ^-1^ at day 56 to 670 mg.kg_sludge_ ^-1^ at day 77 to 0 mg.kg_sludge_ ^-1^ at day 90. The reduction in VFAs led to a rise in pH, restoring neutral conditions and resulting in increased methane production in the initially acidified hydrolysis tank of ABR1 (Fig. 1B, D, F). These findings suggest that OTU_1 plays a role in SPO, consistent with observations for other Cloacimonadota representatives (Westerholm et al., 2022). In contrast, the same compartment of a control ABR0 did not enrich for Cloacimonadota and functioned as a typical hydrolysis tank, with pH oscillating around acidic values (Fig. 1A, C, E; Supplementary Material Fig. S1 – S3). For the purpose of the present study, which focuses on Cloacimonadota and their potential for SPO, we primarily examined and discussed the hydrolysis tank. Results related to the other compartments are provided in the Supplementary Dataset 1 (Table S1-S3) and Supplementary Material (Supplementary Material, Fig. S2 – S3).

To further assess the potential of Cloacimonadota to counteract propionate accumulation, we conducted a bioaugmentation experiment in a third ABR. Its hydrolysis tank, also fed with high OLR of SBP, had become acidified and ceased methane production. To restore its performance, we supplemented it with sludge enriched in the specific Cloacimonadota OTU_1, previously identified in the hydrolysis tank of the test ABR1. After five rounds of bioaugmentation, the acidified compartment resumed methane production, and the pH bounced back to neutral values (Supplementary Material, Results and discussion 2.1). Concurrently, Cloacimonadota OTU_1 established itself in the hydrolysis tank (Supplementary Material, Fig. S5). Therefore, AD sludge enriched in Cloacimonadota OTU_1 can be considered an effective bioaugmentation agent for restoring methane production in acidified AD reactors, as previously recognized in a patent publication (Calusinska et al., 2022).

### Identifying potential syntrophic partners of Cloacimonadota OTU_1

The shift in environmental conditions also altered the archaeal community structure, with species such as *Methanothrix* sp and *Methanospirillum* sp. co-occurring with increased abundance of Cloacimonadota OTU_1 in ABR1 (Supplementary Material, Fig. S5). To investigate whether any of those methanogens could represent potential syntrophic partners of *Cloacimonadota* OTU_1, we performed a CNA based on a 16S rRNA gene amplicon dataset comprising nearly 300 samples from both the present study and previous investigations (Calusinska et al., 2018; Lemaigre et al., 2023). This dataset encompassed a spectrum of anaerobic reactors, ranging from lab-scale to full-scale (Supplementary Dataset 1, Table S2-S3). From this network, *Methanothrix soehngenii* emerged as a keystone taxon, suggesting it serves as the primary archaeal syntrophic partner for Cloacimonadota OTU_1 (Supplementary Material, Fig. S4). Although *Methanothrix sp*. is traditionally classified as an obligate aceticlastic methanogen, genomic analyses have revealed a complete set of genes associated with hydrogenotrophic methanogenesis (Rotaru et al., 2014). Moreover, *Methanothrix* has been shown to participate in direct interspecies electron transfer (DIET), a process that involves interactions with bacterial partners. For example, DIET between *Methanothrix* and *Geobacter* species has been experimentally demonstrated, as summarized by (Lovley, 2017). Other potential syntrophic archaeal partners of Cloacimonadota OTU_1 were taxonomically affiliated with *Methanosarcina* sp. *and Methanospirillum* sp. (Supplementary Material, Results and discussion 2.2; Supplementary Dataset 1, Table S4).

### Genome reconstruction and metabolic insights of *Candidatus* Digestoria delfossei OTU_1: A potential SPO bacterium

Using a combination of short- and long-read sequencing approaches, we reconstructed genomes of 52 microbes residing in the hydrolytic tank of ABR1, comprising eight archaeal and 44 bacterial genomes of medium to high draft genome quality (according to MIMAG standards), mainly representing *Bacteroidota, Spirochaetota, Patescibacteria, Planctomycetota, Desulfobacterota*, and *Halobacterota* phyla (Supplementary Dataset 2, Table S5; Supplementary Material, Fig. S6 and S7). Genome reconstruction of Cloacimonadota OTU_1 yielded a complete, closed genome assembly. It is 2.3 Mbp long, with a GC content of 54.3%, and it contains two rRNA operons (Supplementary Dataset 2, Table S6). The genome was compared to the two described species of Cloacimonadota, *Ca*. Cloacimonas acidaminovorans (Pelletier et al., 2008) and *Ca*. Syntrophosphaera thermopropionivorans (Dyksma and Gallert, 2019) to assess its taxonomic placement. Based on ANI and AAI thresholds for genus and species delineation, it falls below the accepted cutoffs (ANI < 96.5; AAI 60%-80%), indicating it likely represents a new lineage. Adhering to the rules of the Code of Nomenclature of Prokaryotes Described from Sequence Data (SeqCode) for high-quality taxonomic description of uncultivated taxa (Hedlund et al., 2022), we propose naming it *Candidatus* Digestoria delfossei OTU_1 gen. nov., sp. nov. (Supplementary Material, Results and discussion 2.7). Based on genome reconstruction, the *Ca*. Digestoria delfossei encodes a complete glycolysis pathway with exception for pyruvate kinase, previously shown to be replaced by pyruvate orthophosphate dikinase in Cloacimonadota genomes (Johnson & Hug, 2022), and the microorganism can oxidize pyruvate further to acetyl-CoA and acetate. It is capable of beta-oxidation and protein degradation (i.e., encoding multiple peptidases) with a complete metabolism of histidine to glutamate. Although Cloacimonadota has been suggested to represent SPO bacteria (SPOB), a complete *mmc* pathway for propionate oxidation was not found (Supplementary Material, Results and discussion 2.3). The missing steps are as previously reported for this phylum (Westerholm et al., 2022).

### Characterization of encoded metabolic potential of Cloacimonadota

To further evaluate the capacity of Cloacimonadota to oxidize propionate or potentially perform other metabolic roles in the environment, we assembled a collection of Cloacimonadota genomes. This collection included MAGs reconstructed in this and our previous studies and additional MAGs obtained from public databases (Supplementary Dataset 2, Table S7). A phylogenetic reconstruction based on concatenated protein comparisons revealed a grouping of Cloacimonadota into four major clades (I, IIA-C, III, IV), which reflected their environmental origin (Fig. 2). Previously, the existence of two major clades differentiated by ecosystem type was proposed (Johnson & Hug, 2022); our study extends this, including a wider array of ecosystems, as further discussed in Supplementary Material, Results and discussion 2.4 and Fig. S9. Functional annotation identified 2080 KOs, covering an average of 48.3% ± 4.3% of genes per MAG (Supplementary Dataset 2, Tables S8–S9). Over half of the proteins could not be annotated, especially those in termite gut Cloacimonadota, limiting comparative metabolic analysis (Supplementary Material, Results and discussion 2.5; Supplementary Dataset 2, Table S10). Among the functionally assigned genes, clade IIC from marine environments, showed the highest KO diversity (1751 different categories assigned), suggesting broader functional potential than in AD and rock-associated lineages (average 1080 ± 85 KOs). A shared set of 694 Kos likely represents the core Cloacimonadota metabolism, and it is further elaborated in Supplementary Material, Results and discussion 2.5-6 and Supplementary Dataset 2, Table S9 and S11. Comparison between AD and marine, the two environments with the highest number of Cloacimonadota MAGs retrieved in our analysis, indicated 148 KOs enriched in AD MAGs and 109 in marine MAGs (LDA, p < 0.01) (Supplementary Dataset 2, Table S8). Among the KOs enriched in ADs, 107 represented enzymes, including numerous metallopeptidases (e.g., representatives of families M16-M18, M20, M28, M48, and M55) as well as other amino acid related enzymes, lipopolysaccharide biosynthesis proteins and ABC transporters (mainly for sugars). Conversely, in marine MAGs, less than half of the enriched KOs comprised enzymes. Differences in gene enrichment of several mmc pathway genes were observed between marine and AD MAGs, with some genes missing or present in altered forms in marine MAGs (Fig. 3A; Supplementary Dataset 2, Table S8). Notably, several mmc pathway genes, including acetyl transferase, methylmalonyl decarboxylase, methylmalonyl epimerase, oxoglutarate:ferredoxin oxidoreductase, malate dehydrogenase and succinyl-CoA synthetase, were co-encoded in the same genomic location in marine Cloacimonadota, forming a cluster akin to the mmc cluster in other known SPOB (Singh et al., 2023; Westerholm et al., 2022). Additionally, downstream of this putative mmc cluster, marine Cloacimonadota encoded [NiFe] hydrogenase and heterodisulfide reductase. Given the abundance of sulfated organic matter in marine environments, this finding suggests a potential link between SPO and sulfate reduction in this group of Cloacimonadota (Ozuolmez et al., 2020).

**Fig. 3.**
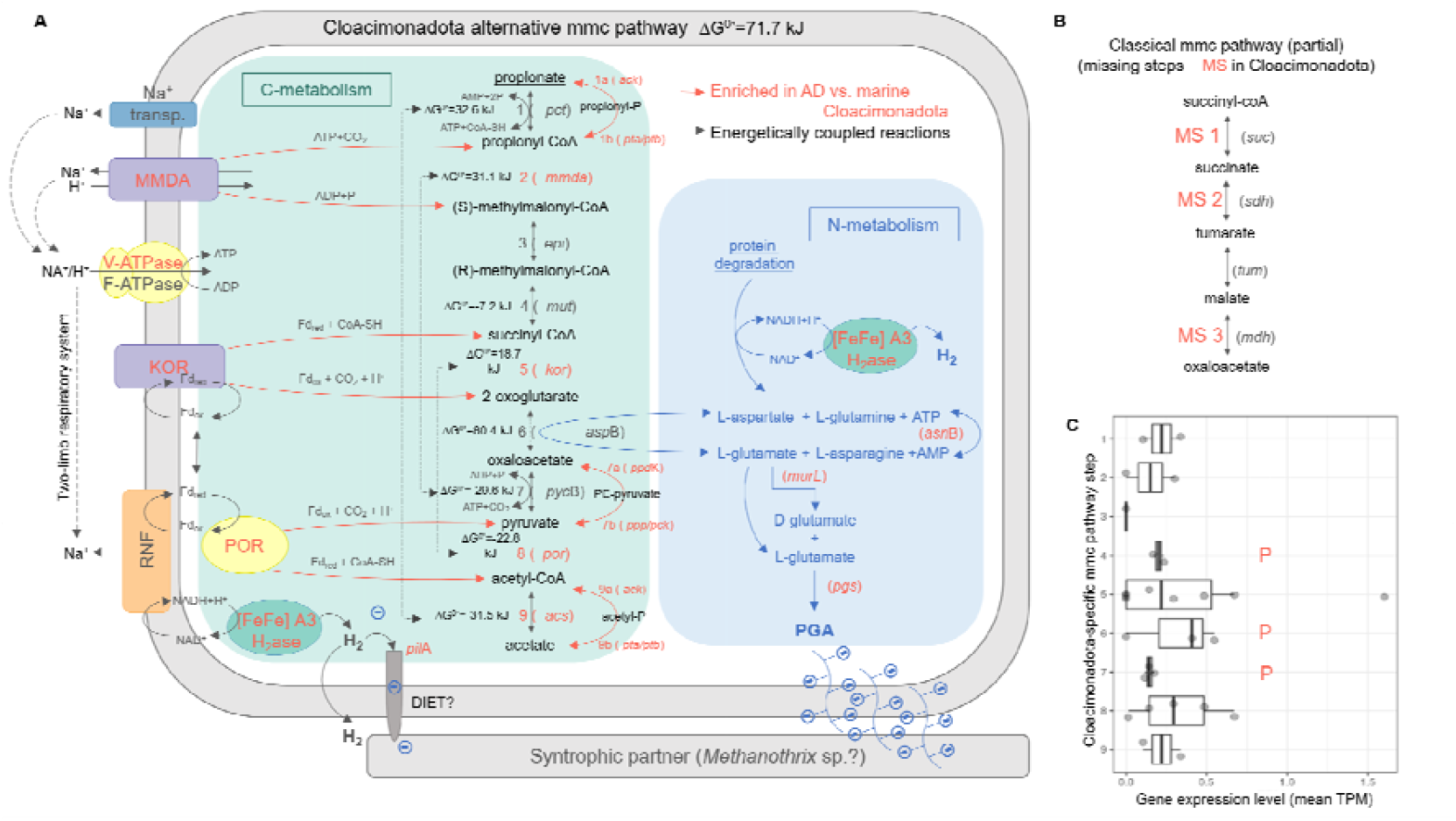
A simplified representation of the proposed syntrophic propionate oxidation pathway in Cloacimonadota found in anaerobic digestion reactors, highlighting its potential connections to nitrogen metabolism and poly-γ-glutamate (PGA) production (A). Missing steps (MS) in the typical mmc pathway (B). Genes and reaction steps overrepresented in AD-associated Cloacimonadota genomes, compared to those from marine habitats, are highlighted in orange. Dashed lines indicate hypothetical links between reaction steps (i.e. energy coupled reactions). Metatranscriptomic abundances of genes involved in different pathway steps are also shown (C). Corresponding proteins identified in the metaproteomic analysis are indicated with the letter ‘P’. Additional details can be found in Tables S13 and S14. MMDA – methylmalonyl-CoA decarboxylase, KOR – 2-oxoglutarate ferredoxin oxidoreductase, POR – pyruvate ferredoxin oxidoreductase, RNF – Rhodobacter nitrogen fixation, [FeFe] A3 H2ase – hydrogenase of group A3, DIET – direct interspecies electron transfer.

### Proposal for an alternative syntrophic propionate oxidation pathway in Cloacimonadota

We hypothesized that SPO is a phylum-wide trait of Cloacimonadota, supported by studies linking propionate degradation to Cloacimonadota OTU abundance in methanogenic reactors (e.g., Dyksma & Gallert, 2019; Nobu et al., 2015). As reported in Westerholm et al. (2022), analysis of two representative Cloacimonadota genomes from methanogenic systems revealed three missing steps of the classical mmc pathway in this phylum (Fig. 3). These include the ATP-generating conversion of succinyl-CoA to succinate catalyzed by succinyl-CoA synthetase (missing step 1), succinate oxidation to fumarate catalyzed by succinate dehydrogenase (missing step 2) and malate oxidation to oxaloacetate catalyzed by malate dehydrogenase (missing step 3). Previous phylum-level analysis of Cloacimonadota genomes also provided no evidence for genes encoding propionate-CoA transferase (Johnson & Hug, 2022), a key enzyme in the conversion of propionate to propionyl-CoA (step 1, Fig. 3A). To explore alternatives for the missing mmc pathway steps in AD Cloacimonadota, we first manually screened the genome of *Ca*. Digestoria delfossei for genes potentially used for propionate oxidation and subsequently verified the presence of these alternative mmc pathway genes in other Cloacimonadota genomes. Based on our analysis, we propose that in Cloacimonadota, the missing steps 1 – 3 of the mmc pathway (Fig. 3B) are substituted first by succinyl-CoA dehydrogenation to 2-oxoglutarate catalyzed by a transmembrane enzymatic complex, 2-oxoglutarate ferredoxin oxidoreductase (kor; Fig. 3A; Supplementary Dataset 3, Table S12 and S13). Subsequently, the generated 2-oxoglutarate is converted by aspartate aminotransferase (aspB) to oxaloacetate, which is further metabolized to acetate via the classical mmc reaction steps. Aspartate aminotransferase facilitates the reversible interconversion of L-aspartate and α-ketoglutarate with oxaloacetate and L-glutamate through a ping-pong catalytic cycle, linking carbohydrate and protein metabolisms (Toney, 2014).

We further screened the other MAGs generated in this study from the ABR reactor to determine whether this alternative mmc pathway might also be present in other species. To our surprise, several *Bacteroidota* MAGs contained at least one gene encoding an enzyme associated with each step of the pathway (Supplementary Material, Results and Discussion 2.3 and Fig. S8), suggesting that the pathway could also be widespread across this phylum. This observation is consistent with previous studies reporting the potential for SPO in diverse bacterial lineages (Hao et al., 2020). However, comprehensive characterization of this pathway in other species is beyond the scope of the present study.

### Comparison of the alternative SPO route with the classical mmc pathway

The thermodynamic calculation of alternative mmc pathway (ΔG^0^’=71.7 kJ; Fig. 3) corroborated the energetic agreement with the classical mmc pathway (ΔG^0^’=73 kJ; Stams & Plugge, 2009). Like the classical mmc pathway, the energy-dependent steps in Cloacimonadota may be coupled with energy-yielding reactions, supported by coupled reactions thermodynamics (Fig. 3A). Cloacimonadota has the potential to connect the initial endergonic step of propionate activation to the last exergonic step of acetyl-CoA deactivation (steps 1 and 9), which is also found in several SPOB. While phosphate butyryltransferase (K00634) seems prevalent in Cloacimonadota genomes (Supplementary Dataset 2, Table S8), butyrate oxidation has not been reported by members in this phylum so far. This suggests that this enzyme could have been misidentified as a phosphate propionyl transferase (encoded by the *pta* gene; Fig 3A). Indeed, information from the Brenda database suggests that the same enzyme can accept propionyl-CoA alongside butyryl-CoA in *Listeria monocytogenes* (Sirobhushanam et al., 2017). Interestingly, compared to Cloacimonadota found in AD reactors, marine Cloacimonadota exhibit gene enrichment in an alternative enzyme known as acetyl-CoA synthetase (*acs*). This suggests that the propionate activation step can also be driven autonomously. Step 2 in propionate degradation in Cloacimonadota entails a membrane-bound Na^+^ - transporting methylmalonyl decarboxylase (*mmda*), coupled with oxaloacetate decarboxylation to pyruvate (step 7). Methylmalonyl decarboxylase relies strictly on Na^+^ ions for activity. Transportation across the cytoplasmic membrane creates a sodium ion motive force, utilized for ATP synthesis. These enzymes, along with the putative phosphate (butyrate) propionyltransferase (*pta*, step 1) and methylmalonyl-CoA epimerase (*epi*, step 3) are among the most frequently encountered protein clusters in the Cloacimonadota gemones (Supplementary Dataset 2, Table S10; Supplementary Material, Fig. S10). Their high conservancy suggests that they play a fundamental role in the metabolism of this phylum. Different forms of methylmalonyl-CoA mutase (*mut*), an enzyme catalyzing step 4 in the propionate degradation pathway, are present in both AD and marine Cloacimonadota. K01847 is enriched in AD, while the dimeric form (K01848 and K01849) is found in marine genomes. The transmembrane 2-oxoglutarate:ferredoxin oxidoreductase (*kor*) is encoded in the genomes next to the methylmalonyl-CoA epimerase, likely forming a single operon, which further supports its role in the propionate degradation pathway in Cloacimonadota. This reaction (step 5) is likely coupled to one catalyzed by pyruvate:ferredoxin oxidoreductase (*por*, step 8). In the absence of a ferredoxin-dependent hydrogenase in Cloacimonadota (Losey et al., 2020), this enzyme would recycle electrons from a reduced ferredoxin, which would otherwise be used for H_2_ production in other species (Calusinska et al., 2010). The conversion of 2-oxoglutarate to oxaloacetate is putatively linked to the transamination of L-aspartate and L-glutamate (step 6). This step would connect carbohydrate and protein metabolisms, with 2-oxoglutarate acting as a major carbon skeleton to assimilate nitrogen from protein degradation. Indeed, the ability of Cloacimonadota to ferment amino acids to 2-oxoacids for carbon and energy has previously been reported (Dyksma & Gallert, 2019; Pelletier et al., 2008). Based on the presence of genes encoding additional enzymes involved in the interconversion/transformation of glutamate ↔ glutamine (i.e., glutamate dehydrogenase, glutamine synthase) and aspartate ↔ asparagine (asparaginase) in the genomes of nearly all AD Cloacimonadota, we further hypothesize that Cloacimonadota could exclusively feed on propionate if specific N-precursors are present in the medium.

RNA sequencing (metatranscriptomics) revealed the transcription of all genes identified as part of the proposed alternative SPO pathway (Supplementary Dataset 3, Table S14), except for methylmalonyl-CoA epimerase (*epi*, step 3; Fig. 3C). However, the read counts for these genes were relatively low (Supplementary Material, Fig. S7). Metaproteomic analysis further identified several proteins representing various steps of the proposed pathway under high propionate conditions, including PTA (step 1b), MUT (step 4), ASPB (step 6), PYC B (step 7), as well as components of RNF, trimeric A3 [FeFe] hydrogenase, and V-ATPase complexes (Supplementary Dataset 3, Table S15). Additionally, glutamate dehydrogenase and L-asparaginase were abundant, indicating active pathways involved in nitrogen metabolism linked via step 6 to the proposed SPO pathway. Collectively, these findings provide tentative support for the validity of the newly proposed SPO pathway in Cloacimonadota.

### Distinct metabolic role of Cloacimonadota compared to other bacteria present in the anaerobic digestion environment

We further aimed to specifically characterize the distinct role of Cloacimonadota within the AD microbiome, with a focus on evaluating their phylum-wise metabolic capacity for SPO (Supplementary Dataset 4, Table S16 and S17). To achieve this, we utilized another extended collection of previously published AD-specific MAGs (Campanaro et al., 2020), which we supplemented with Cloacimonadota from our database. Genome-scale analysis, including gene assignment to Pfam domains and NMDS projection of Jaccard distances, revealed some well-defined phylum-level clusters on the NMDS graph, with the *Cloacimonadota* cluster notably distinct (Fig. 4A). Reasoning that ubiquity indicates essential gene functions, while abundance without wide distribution across various phyla suggests adaptive, habitat-specific functionalities, we initially focused on functions (i.e. KOs) predominantly present in Cloacimonadota (Fig. 4B). Acknowledging the incompleteness of some microbial genomes used in our study, we established criteria for functions represented in at least 80% of Cloacimonadota genomes and absent in at least 80% of other bacterial genomes commonly found in AD environments. As a result, we found only six KOs that were ubiquitous in all AD bacterial genomes but nearly absent from *Ca*. Cloacimonadota. These KOs were mainly involved in cell biogenesis and phosphate starvation inducible stress (K06217). The latter enables cells to use limited phosphate resources more effectively (Antelmann et al., 2000).

**Fig. 4.**
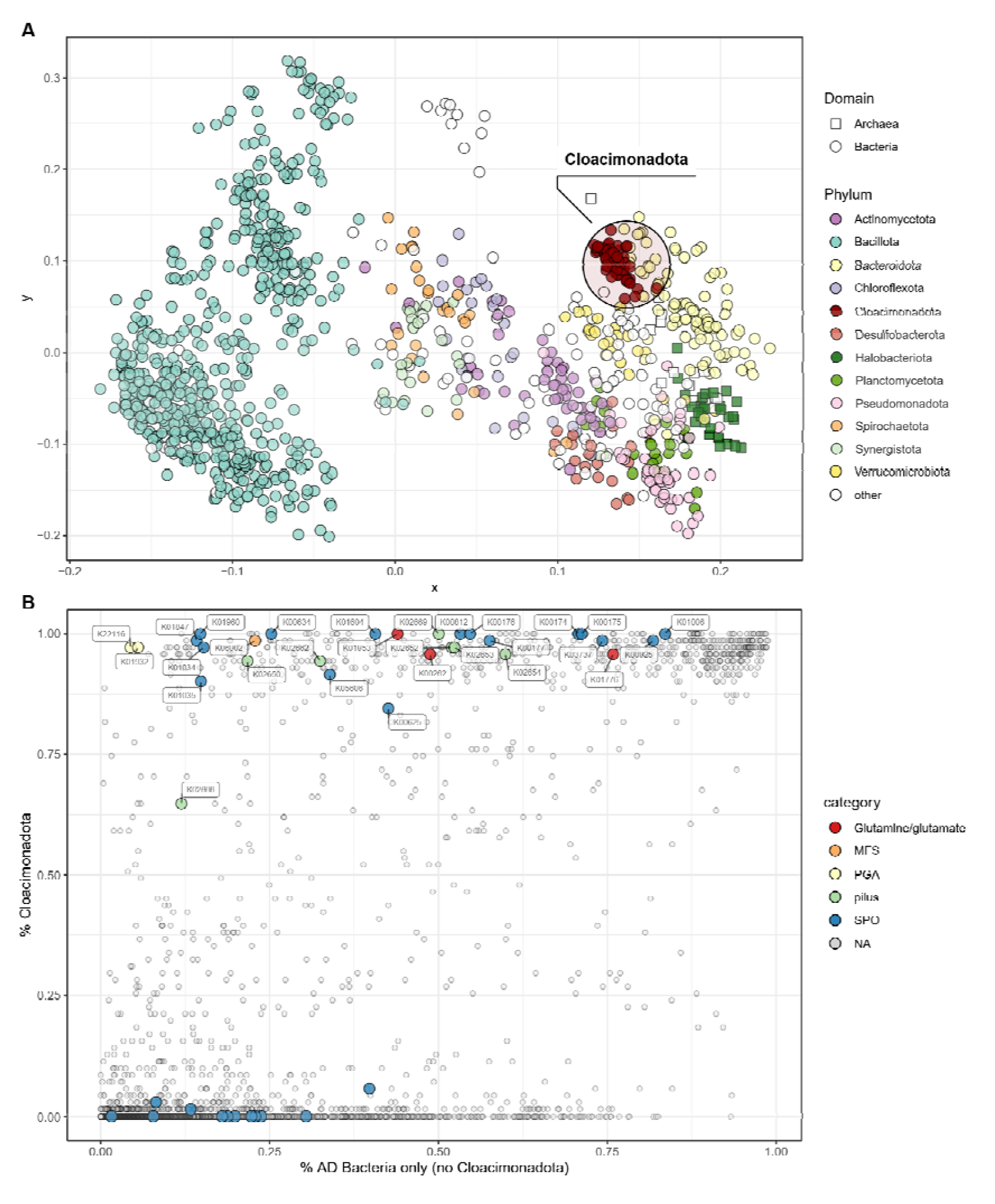
Visual representation of the specific functions of Cloacimonadota within the AD microbiome. Non-metric multidimensional scaling (NMDS) of Pfam domains present in Cloacimonadota genomes compared to other AD phyla (A). Overrepresentation of KEGG Orthologies (KOs) in Cloacimonadota genomes compared to other AD bacteria (B). KOs involved in syntrophic propionate degradation and associated pathways (referenced in Fig. 3) are highlighted in the graph. MFS – Major Facilitator Superfamily transporter, PGA – poly-γ-glutamate, SPO – syntrophic propionate oxidation.

Conversely, we identified 58 KOs (39 with enzymatic function) nearly exclusively assigned to Cloacimonadota (Supplementary Dataset 4, Table S16), including those related to sodium (K09696, K09697) and phosphonates (K02041, K02042) ABC transport, lysin degradation (K01844, K18011, K18012, K18013), various metallo- (K01284, K06972, K01267, K05823, K05994) and serine-peptidases (K03503, K08676), multiple uncharacterized proteins (K09129, K09797, K09141, K09798, K09729) and KOs involved in lipopolysaccharide biosynthesis (K16363, K03269, K09778, K03270). The specialization of Cloacimonadota towards the plausible utilization of phosphonates, organophorous compounds with C-P bonds, as an alternative phosphorous source, might indicate that this phylum has carved out a specific niche within the AD community by utilizing a resource that is largely inaccessible to other organisms (Sosa et al., 2017). Indeed, sugar beet pulp has high fiber and protein content but low phosphorus content, which could stimulate the overdominance of Cloacimonadota in our test reactor (Fig. 1). However, this observation remains to be further studied. Among other KOs widely present in Cloacimonadota genomes, four KOs, including methylmalonyl-CoA mutase (K01847), acetate CoA transferase (K01035, K01034) and pyruvate carboxylase (K01960) are commonly associated with the mmc pathway in known SPOB (Fig. 4B). All other mmc genes were slightly below the threshold, indicating they were also frequently detected in other AD phyla, yet they were still present in over 90% of the AD Cloacimonadota genomes (Fig. 4B). By analogy to a recently characterized major facilitator superfamily (MFS) transporter responsible for propionate tolerance in a Gram-negative bacterium *Pseudomonas putida* (Ma et al., 2021), we identified a K06902 as a putative propionate transporter gene in Cloacimonadota. Its presence in nearly 99% of analyzed Cloacimonadota genomes, while roughly 22% of other AD bacteria also encoded a respective gene, further reinforce a rare metabolic function of Cloacimonadota in SPO among AD microbes. The L-glutamate-generating aspartate aminotransferase (K00812), which we hypothesize compensates for the absence of certain steps in the classical mmc pathway, was prevalent in most of the analyzed Cloacimonadota genomes, sometimes occurring in multiple copies. In contrast, only half of the other AD microbes encoded the respective gene(s). Interestingly, the near-ubiquitous presence of a PGA biosynthesis (K01932, K22116) cluster in AD Cloacimonadota (Supplementary Material, Fig. S12), compared to its rarity in other AD bacteria (identified in less than 5% of genomes, mainly *Bacillota*, formerly *Firmicutes*), suggests a potential role for L-glutamate in PGA production (see below). Additionally, aspartyl aminopeptidase (K01267), commonly found in Cloacimonadota but relatively rare in other AD microbes, can release N-terminal aspartate and glutamate from pre-digested peptides, facilitating their entry into the cycle.

### Association of poly-γ-glutamate production with propionate degradation in Cloacimonadota

Building on our previous observation that the SPO may be coupled with the poly-γ-glutamate formation, we examined this potential link more closely in the recovered Cloacimonadota genomes. The PGA operon of *Ca*. Digestoria delfossei OTU_1 is composed of three genes encoding proteins similar to CapB, CapC and PgsF, and exhibits the typical gene organization found in know Gram-negative PGA producers (Supplementary Material, Fig. S12). PGA production involves L-glutamic acids units derived exogenously or endogenously using 2-oxoglutaric acid as a precursor (Fig. 3A). In the absence of a complete TCA cycle in Cloacimonadota, we hypothesize that 2-oxoglutaric acid derived from the succinyl-CoA carboxylation mediated by the 2-oxoglutarate:ferredoxin oxidoreductase, could directly link PGA production to propionate degradation (Fig. 3). Nevertheless, its production from exogenously provided L-glutamic acids units is also possible, as we identified a putative glutamate ABC transporter in the genome of *Ca*. D. delfossei. The respective operon was also identified in 98% of analyzed Cloacimonadota genomes, further suggesting a phylum-wise capacity for glutamate transport.

Overall, poly-γ-glutamic acid production has been primarily studied in Bacillus species, where the polymer was initially shown to be involved in virulence (i.e. antibiotic resistance), such as in *Bacillus anthracis* (Scorpio et al., 2007). However, the presence of a capsule synthesis protein domain “IPR019079” named CapA, which is involved in PGA production, has also been observed in short chain fatty acids degrading syntrophic bacteria (Worm et al., 2014). Primarily, Gram-positive bacteria are known to produce PGA, while only a few Gram-negative bacteria have been identified to synthesize this polymer (e.g. Candela & Fouet, 2006). The physiological function of PGA in different organisms is not fully understood and depends on whether the polymer is anchored (i.e. a virulence factor) to the peptidoglycan or released (Luo et al., 2016). In the latter case, it can provide environmental advantages, such as helping to sequester toxic metals, decreasing salt concentration, providing a carbon source, protecting against adverse conditions, and improving biofilm formation. We speculate that in Cloacimonadota, PGA might promote syntrophic interactions by facilitating direct contact with its syntrophic partner(s), thus enhancing SPO. Moreover, highly charged polyanions such as PGA exhibit electrical conductivity behavior (Bizzarri et al., 1990), which could potentially facilitate DIET between syntrophic partners. DIET has been previously suggested for Cloacimonadota (summarized in Westerholm et al., 2022), and we further identified type IV pilus assembly proteins in almost all AD Claocimonadota genomes. In *Ca*. D. delfossei, the entire operon was identified and is encoded directly upstream of some of the mmc pathway genes (i.e. the cluster containing the methylmalonyl epimerase and 2-oxoglutarate oxidoreductase genes; Supplementary Material, Fig. S11), indicating potential common regulation mechanisms.

### Culture enrichment of *Ca.* Cloacimonas fortuita, another species of Cloacimonadota from anaerobic digesters

Several genome-informed cultivation trials were conducted using different anaerobic media and carbon sources to selectively enrich Cloacimonadota, with the aim of isolating *Ca*. Digestoria delfossei OTU_1 (Fig. 5; Supplementary Material, Results and discussion 2.7). Despite multiple attempts, we were unable to isolate this species. However, in ampicillin and streptomycin fortified DMS 621 medium using D-glucose as a carbon source (culture named “RT_BT_as_ph7”), we managed to enrich another Cloacimonadota species, which was subsequently passaged on a medium containing propionate as the only carbon source (Supplementary Dataset 5, Table S18 – S21). We attribute its antibiotic resistance to the presence of the PGA protective matrix surrounding the cells, given that no specific antibiotic resistance was identified in the Cloacimonadota genomes (data not shown). The culture was rather slowly growing, forming loose white flocks (Fig. 5), which were easily destroyed if the bottle was vigorously shaken. Based on 16S rRNA gene amplicon sequencing, the corresponding OTU accounted for 56% of the community, with only two other bacterial OTUs showing significant abundance (Supplementary Dataset 5, Table S19). We were pleased to discover that the archaeon also preset in the enriched culture was actually *Methanothrix soehngenii* (Supplementary Dataset 5, Table S21), initially identified as a putative partner of *Ca*. D. delfossei based on the CNA (see paragraph “Identifying potential syntrophic partners of Cloacimonadota OTU_1”). Simple microscopy observation of the Gram-stained culture visualized the negatively stained sphere-shaped Cloacimonadota-like cells and long filaments typical of *Methanothrix* being embedded in a matrix, possibly composed of PGA (Fig. 5C). Sporadically, coccoidal aggregates typical to *Methanosarcina* genus (another putative partner based on the CNA), were also present (not shown), and fragmented genome pieces were re-constructed from metagenomics reads as well (Supplementary Dataset 5, Table S21). FISH analysis confirmed the presence of Cloacimonadota conglomerates, densely embedded in the PGA (Fig. 5), as could be visualized by DAPI staining of PGA based on (Szczepanek et al., 2002). The draft genome of this new *Ca*. Cloacimonadota species is 1.98 Mbp long, with a GC content of 36.6%, and it contains 1529 protein coding genes (Supplementary Dataset 2, Table S6). Comparing it to the two others described Cloacimonadota genomes, e.g. *Ca*. Cloacimonas acidaminovorans and *Ca*. Syntrophosphaera thermopropionivorans, shows that it represents a new species (Supplementary Material, Fig. S13) within the Cloacimonas genus, hereinafter named *Candidatus* Cloacimonas fortuita RT_BT_as_ph7 sp. nov. (Supplementary Material, Results and discussion 2.7).

**Fig. 5.**
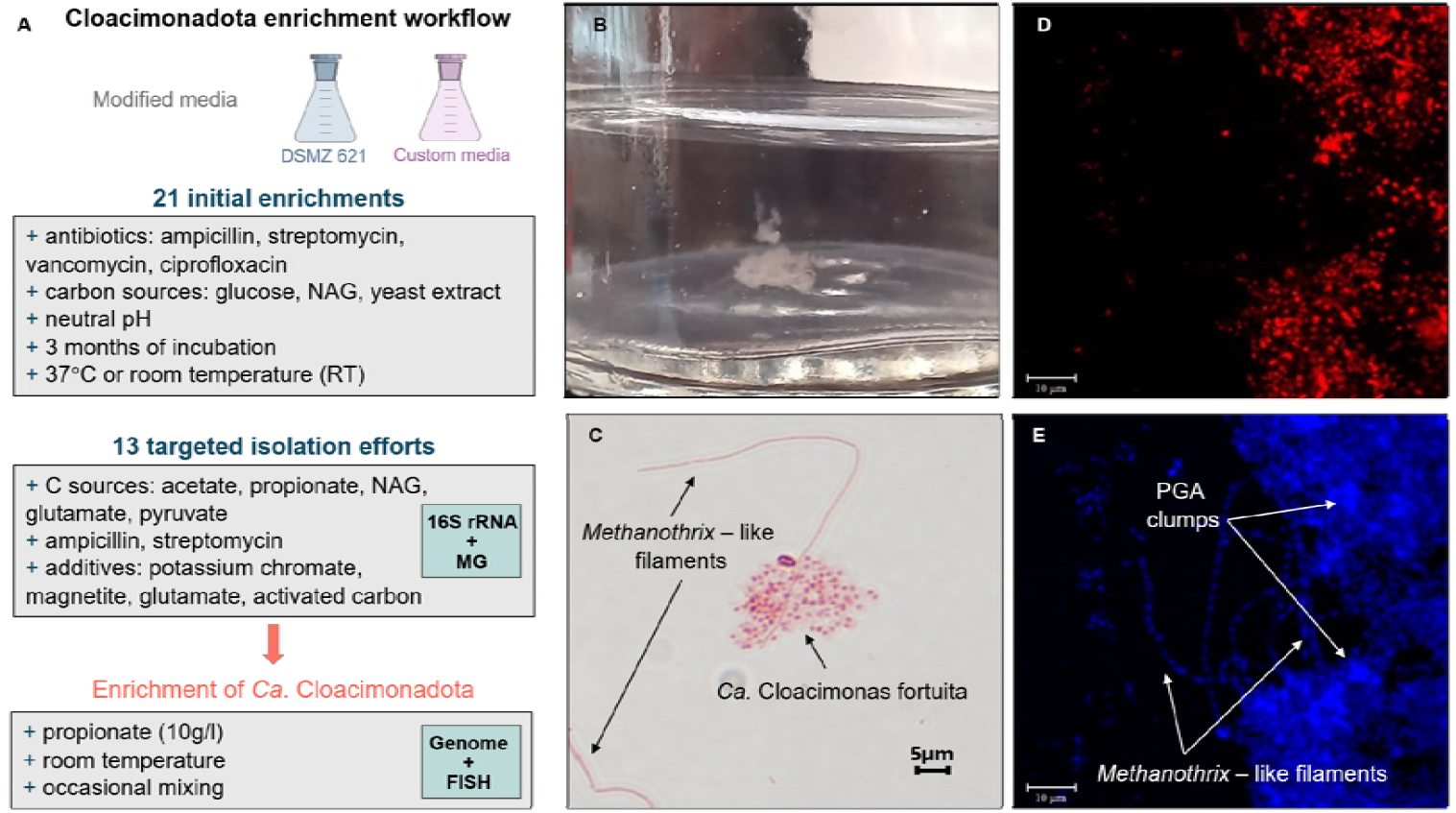
Enrichment assay design (A) and microscopy results of cultures enriched with *Ca*. Cloacimonas fortuita. Formation of loosely aggregated flocs (B). Gram staining (C). Fluorescence in situ hybridization (FISH) images showing the presence of enriched Cloacimonadota species in red (D). DAPI staining of the enriched culture highlighting the association of *Methanothrix* sp. with Cloacimonas fortuita and the formation of poly-γ-glutamate (PGA) clumps (E; following the PGA-DAPI staining method described by Szczepanek et al., 2002).

## Conclusions

Our study highlights the ecological importance of Cloacimonadota in methanogenic environments and underlines that syntrophic propionate oxidation could be a conserved functional trait within this bacterial lineage. Cloacimonadota are consistently abundant in methanogenic reactors that exhibit propionate accumulation, suggesting a strong ecological link to propionate turnover under anaerobic conditions. Genomic analysis further reveals that they encode a complete set of genes for SPO, though not following the classical mmc route. Instead, they utilize an alternative enzymatic route that is thermodynamically equivalent to the classical mmc pathway. Both sequencing results and microscopic observations revealed that Cloacimonadota co-occur with methanogenic archaea, strongly suggesting close interactions. Network analysis and Cloacimonadota enrichments point to *Methanothrix* as a key archaeal partner in the syntrophic cooperation. Still, further research is needed to determine the conditions under which this partnership is established and how it functions. A key limitation of this study is the inability to isolate and maintain Cloacimonadota in pure culture. Although some species can be studied within enrichment communities, as evidenced by the isolates in our study namely *Ca*. Digestoria delfossei and *Ca*. Cloacimonas fortuita, their growth is unstable and further advancements in cultivation methods will be essential. As such, future efforts should aim at isolating Cloacimonadota representatives in defined co-culture with the archaeal partners to experimentally validate their metabolic interactions. Furthermore, elucidating the regulatory mechanisms governing the SPO and PGA-based biofilm formation may offer deeper insights into the ecological success of Cloacimonadota in the propionate-intoxicated methanogenic environments.

## Acknowledgements

We thank Bénédicte de Vos and Tommaso Serchi from LIST for their valuable technical support with reactor operations and microscopy.

## Data availability

The 16S rRNA gene sequences generated in this study along with the metagenomic and metatranscriptomic datasets are deposited in the NCBI BioProject accession number PRJNA1320513. The 16S rRNA gene sequences used in this study were previously reported in our earlier works and can be accessed through the cited references (Calusinska et al., 2018; Lemaigre et al., 2023). The genomes of newly enriched Cloacimonadota species have been deposited in SeqCode: the complete and closed genome of *Ca*. Digestoria delfossei is available in registry https://seqco.de/i:52909, and the high-quality draft genome of *Ca*. Cloacimonas fortuita is available in registry https://seqco.de/i:52907.

