## Supplementary Material for "Phylum-wide propionate degradation and its potential connection to poly-γ-glutamate biosynthesis in *Candidatus* Cloacimonadota phylum"

**Supplementary Methods**

1.1 **Anaerobic baffled reactors and experimental design**

**Operation of reactors**

Two three-compartment anaerobic laboratory-scale baffled reactors (ABRs) with a capacity of 100 L were inoculated with anaerobic sludge from a full-scale methanogenic reactor at the wastewater treatment (WWTP) plant in Schifflange, Luxembourg. The ABRs operated at 37°C over a period of 174 days (Fig. 1), each reactor (equal size, 33L volume) was compartmentalized by two baffles, with the first compartment (i.e., hydrolysis tank) receiving feed. The ABR reactors were manually fed with commercial dried sugar beet pulp (SBP) pellets following a semi-continuous feeding scheme, receiving feed on working days only, with no feeding on weekends (Fig. S1). Reactors were fed at an organic loading rate (OLR) increasing from 0.5 to 3 kg VS m^-3^d^-1^ for ABR0 (control reactor; Fig. 1A, C, E) and an OLR of 0.5 to 6 kg VS m^-3^d^-1^ for ABR1 (test reactor, Fig. 1B, D, F). The pulp contained 0.012 grams of nitrogen (N) per gram of volatile solids (g_N_.g_VS_^-1^), and its biochemical methane potential (BMP) was measured to be 0.38 normalized liters of CH_4_ per gram of volatile solids (VS; LN_CH4_.g_VS_^-1^), according to (Lemaigre et al., 2023). The hydraulic retention time (HRT) was adjusted to 56 days by adding tap water to the SBP pellet. A total of 7L of sludge was re-circulated from the last compartment to the first, i.e., the hydrolysis tank, each time the reactors received feed (Fig. S1). Samples were regularly collected from each reactor compartment for analytical analyses. Metadata, including gas composition, pH, total ammonium nitrogen (TAN; g_N L_sludge_^-1^), total inorganic carbon (TIC; g_CaCO3 L_sludge_^-1^), total solids (TS; % of total solid/fresh sludge [w/w]), volatile solids (VS; % of volatile solids/% of total solids [w/w]), and volatile fatty acids (VFAs; g.kg_sludge_^-1^), were analyzed according to (Lemaigre et al., 2023).

Biogas production was measured automatically every two hours, as previously described by (Lemaigre et al., 2023). Methane content in the biogas was estimated using gas chromatography (CompactGC, Global Analyser Solutions, Interscience). Volatile fatty acids (VFAs), including formate, acetate, propionate, iso-butyrate, n-butyrate, and caproate, were monitored as described below. Briefly, approximately 250 μL of collected digestate was centrifuged at 15,000 x g for 5 minutes at 4°C. A total of 150 μL of the supernatant was collected and mixed with 600 μL of distilled water. Samples were filtered through PVDF 0.45 μm filters, and 50 μL of the collected filtrate was mixed with distilled water. Samples were stored at -20°C if not analyzed immediately. VFAs were measured by ion-exchange chromatography with conductivity detection and electrochemical suppression using an Ion Chromatograph ICS 5000 Dual-Channel from Thermo Fisher Scientific. VFAs were eluted with KOH on a Dionex IonPac AS18 column equipped with a guard column AG18.

**Bioaugmentation trials**

In addition to the control (ABR0) and test (ABR1) reactors, a third ABR (ABR2), also fed with SBP, was used in the bioaugmentation trials (Fig. S5). The aim of this experiment was to further evaluate the potential application of the reactor microbiome enriched with Cloacimonadota OTU_1 as a remedy for treating acidified reactors with significant propionate concentrations. The fed compartment (hydrolysis tank) of the bioaugmented ABR2 was highly acidified, with a pH below 5.0, propionate concentrations above 5000 mg.kg^-1^ of sludge, and methane content in the produced biogas of less than 30% (Supplementary Dataset 1, Table S1). The bioaugmentation procedure began on day 132 and continued for 5 weeks. In total, 5 bioaugmentation events were applied, during which 3 L of sludge from the fed compartment of the test reactor (ABR1) were transferred to the first compartment (hydrolysis tank) of the bioaugmented reactor (ABR2).

**1.2 16S rRNA gene amplicon sequencing and correlation network analysis**

**Library preparation, sequencing, and data analysis**

Macromolecules DNA/RNA were co-extracted using the Allprep DNA/RNA Mini kit (Qiagen, Hilden, Germany), following the manufacturer's instructions. The eluate was then divided into two parts: one part was treated with 1 μL of 10 μg/ml RNaseA (Sigma) for 30 minutes at room temperature, and the other part was treated with TURBO DNase (Invitrogen) according to the manufacturer's instructions, to obtain pure DNA and RNA fractions, respectively. The quality and quantity of extracted nucleic acids were assessed using the Bioanalyzer (Agilent) and Qubit (Invitrogen), respectively. DNA extracts were stored at -20°C until further library preparations. RNA extracts were stored at -80°C until further library preparations.

The bacterial and archaeal 16S rRNA gene amplicon libraries were sequenced using the Illumina MiSeq approach, as previously described (Calusinska et al., 2018; Goux et al., 2015). In summary, a modified version of universal bacterial primers S-D-Bact-0909-a-S-18 and S--Univ--1392-a-A-15, and archaeal primers S-D-Arch-0519-a-S-15 and S-D-Arch-1041-a-A-18 (Klindworth et al., 2013), together with the Nextera XT Index Kit V2 (Illumina), were used in a two-step PCR reaction. This amplified a fragment of approximately 484 bp, spanning the V6–V8 region of the bacterial 16S rRNA gene, and a fragment of around 526 bp, spanning the V4-V6 region of the archaeal 16S rRNA gene. Following sequencing, Usearch v.7.0.1090_win64 software was used for quality trimming (fastq-maxee 1, fastq_minlen 400 for bacteria and fastq_minlen 500 for archaea), chimera checking, removal of singletons, and assignment of sequences to operational taxonomic units (OTUs) at the 97% similarity level, according to the pipeline described previously (Edgar, 2010). The taxonomic affiliation of the resulting OTUs was performed using the SILVA database v.138.2 (Yilmaz et al., 2014) with mothur (v.1.38.0; Schloss *et al.*, 2009). The sequencing reads are available in the Sequence Read Archive (SRA) database under bioprojectID PRJNA1320513.

**16S rRNA amplicon datasets for correlation network calculation**

We performed correlation network analysis (CNA) using two distinct datasets containing sequences corresponding to our *Cloacimonadota* OTU_1. The first dataset comprised sequences analyzed in this study, combined with amplicon sequence variant (ASV)-level data published by (Lemaigre et al., 2018). The second dataset included a year-long monitoring dataset from (Calusinska et al., 2018), as originally published at the OTU level. CNA was applied separately to each dataset, and the results were compared afterward. For both datasets, bacterial and archaeal sequences were incorporated into the analysis to capture potential interactions across the microbial community.

**Correlation Network Analysis**

The method for calculating pairwise correlations and data filtration was adapted from (De Vrieze et al., 2018). To enhance the sensitivity of resulting networks, infrequent OTUs/ASVs were filtered out, retaining only those representing at least 0.1% of the total community abundance, as recommended by (Berry & Widder, 2014). Subsequently, pairwise Spearman’s rank correlations were computed, and the resulting correlation *p*-values were corrected for multiple comparisons using the Benjamini-Hochberg correction. Data filtering retained only highly significant correlations, defined as those with a *p*-value ≤ 0.001 and an R coefficient ≥ 0.5. Initially, general correlation networks were constructed. To identify potential syntrophic partners of Cloacimonadota OTU_1, direct neighborhood correlation networks were then constructed using only OTUs/ASVs highly positively or negatively correlated with our Cloacimonadota species of interest. Topological features such as Degree Centrality, Betweenness Centrality, Closeness Centrality, and Eigenvector Centrality were calculated for each microbial correlation network to identify potential key actors (*i.e.,* keystone OTUs/ASVs; Supplementary Dataset 1, Table S4). All calculations were performed using R version 3.6.1, RStudio version 1.1.383, Cytoscape version 3.7.2, along with the R Bioconductor package RCy3 version 2.4.4 and the R package igraph version 1.2.4.1.

**1.3 Metaproteomics**

Two samples (Fig. 1), including technical duplicates, were centrifuged at 4°C at 10 000 g during 20 min in an Allegra 64R centrifuge (Beckman Coulter, USA). The pellet was suspended in 600 µl of SDS buffer (30% sucrose, 2% sodium dodecyl sulphate (SDS), 0.1 M Tris-hydrochloride (HCl), 5% β-mercapto-ethanol, pH = 8), vortexed and incubated at 65 °C for 1 h 30. After incubation, the tube was filled with phenol buffer (Invitrogen, Thermofisher scientific) at room temperature, vortexed for 30 s and centrifuged at 10,000 *g* for 3 min. Three phases (lower phase = phenol + cellular debris, upper phase = aqueous SDS phase with solubilized proteins and fat) were formed in the tube and the upper phase was extracted into a new tube. The extract was diluted in cold 0.1 M ammonium acetate in methanol and kept at −20 °C for the night, after which it was centrifuged at 10,000 *g* for 5 min. The pellet was washed twice with cold acetone and dried. Finally, the dried pellet was solubilised in 50 µl lysis buffer (7 M urea, 2 M thiourea, 0.5% (*w*/*v*) CHAPS). Protein concentration was determined following the RC DC ^TM^ (reducing agent, detergent compatible) protein assay (Bio-Rad) with bovine serum albumin (BSA) for the standard curve. Samples were kept at −20 °C until further analysis.

20 µg of total proteins were loaded and separated on a Criterion ^TM^ XT precast 1D-gel (4–12% bis-tris, 1.0 mm × 12 wells, Bio-Rad, USA) following manufacturer's instructions. After a short migration, gels were stained, cut into small pieces for each sample to perform in-gel digestion. Each sample was reduced, alkylated and destained. Then, proteins were digested using trypsin enzyme (sequencing mass grade, Promega, USA). The extracted peptides were analysed with a NanoLC 425 Eksigent coupled to a TripleTOF® 6600 MS (Sciex, Belgium). Peptides were loaded onto the trap column (C18 acclaim™ PepMap™, 5 µm, 5 mm × 300 µm, Thermo Scientific, Germany) and desalted for 5 min at a flow rate of 2 µl/min using loading buffer (2% v/v acetonitrile, 0.05% (v/v) trifluoroacetic acid in water LC-MS grade). After this, peptides were separated onto a C18 reverse phase column at a flow rate of 300 nl/min (C18 acclaim™ PepMap™ 100, 3 µm, 100 A, 75 μm × 15 cm, Thermo Scientific, Bremen, Germany) using a binary gradient (solvent A: H_2_O LC-MS, 0.1% (v/v) formic acid; solvent B: acetonitrile, 0.1% (v/v) formic acid). Peptides were eluted from 3% B to 30% over 60 min, increased to 40% B during 10 min then increased to 80% B until 10 min, and then re-equilibrated prior to the next injection for 20 min at 3% B. MS scan was followed by 30 MS/MS scans from 300 to 1250 m/z with 250 ms of accumulation time and from mass range 100–1500 m/z with 50 ms of accumulation time respectively using the automatically adjusted system of rolling collision energy voltage.

The acquired MS and MS/MS data were imported into Progenesis QI for Proteomics software (version 4.2, Nonlinear Dynamics, Waters). Then the protein and peptide identification were imported to Progenesis QIP, searching against *Ca.* Digestoria delfossei genome via Mascot Daemon (version2.6.0. Matrix Science, UK) and matched to peptide spectra. The following Mascot research parameters were used: peptide tolerance of 20 ppm, fragment mass tolerance of 0.5 Da, a maximum of two missed cleavages, carbamido-methylation of cysteine as fixed modification and oxidation of methionine, N-terminal protein acetylation and tryptophan to kynurenine as variable modifications. Only the proteins identified with a significance Mascot-calculated confidence of 95% and at least two sequences and one unique sequence per protein were accepted. Results are provided in Supplementary Dataset 3, Table S15.

**1.4 Enrichment and isolation trials**

The initial enrichment of bacteria from anaerobic digestion (AD) systems was part of a larger study investigating microbial communities in these environments (details not described here). Two inocula were used, including thickened material from AD systems fed with either activated sludge or agricultural biowaste. Before media inoculation, the sludge inocula were pre-treated with broad-spectrum antibiotics (see below) and diluted 10- or 100-fold. From these enrichments, 21 promising cultures that retained the presence of Cloacimonadota were selected for further tailored cultivation. To promote the growth of Cloacimonadota, various media and cultivation conditions were tested, at neutral pH and incubation temperatures of room temperature or 37°C (Supplementary Dataset 5). Given that Cloacimonadota are hypothesized to produce extracellular polymeric substances composed of poly-gamma-glutamate, we speculated that this feature could enhance resilience to environmental stressors by acting as a protective layer, including against antibiotics. As a result, despite their putative sensitivity to certain antimicrobials (data not shown), a combination of broad-spectrum antibiotics (ampicillin, ciprofloxacin, streptomycin, and vancomycin) was used to suppress the growth of competing bacteria (Supplementary Dataset 5, Table S18). At the end of the incubation period, DNA was extracted from the samples for microbial community analysis using 16S rRNA gene amplicon sequencing and metagenomics.

**1.5 Fluorescence in situ hybridization and microscopy**

Flock formation was observed within the culture media, which could be disrupted by vigorous shaking. To enable microscopic visualization of physical associations between Cloacimonadota and its putative syntrophic partner *Methanothrix*, culture samples were handled gently to minimize disturbance and avoid centrifugation steps. Floating flocks were harvested using a sterile syringe and needle, fixed in 4% paraformaldehyde (w/v) at a 1:3 ratio, and incubated for 3 hours at 4°C. After fixation, samples were washed twice in PBS buffer and stored in a PBS (1:1) solution at -20°C until further processing. Between 5 to 10 μl of sample was applied onto a 10-well FISH slide and air-dried overnight. Subsequently, slides were dehydrated using a series of ethanol solutions with concentrations of 50%, 80%, and 100% for 3 minutes each, followed by air-drying. The hybridization buffer with a 40% formamide concentration was pre-warmed to the desired temperature and applied to the slide. Probes were then added at a final concentration of 4.5 pmol.μl^-1^. Slides were placed in a 50-mL Falcon tube containing moistened paper and incubated horizontally in a hybridization oven for 2 hours at 46°C. After hybridization, slides were washed with the wash buffer at 48°C for 25 minutes. For microscopy, a DAPI-containing antifade mounting solution (Fluoroshield with DAPI, Merck) was applied to the slide and covered with a coverslip. Visualization was performed using a Zeiss LSM 880 system coupled with an inverted Zeiss Axiovert 200 M microscope (Zeiss, Jena, Germany) for image acquisition. Image processing and visualization were conducted using Zeiss ZEN 2011 software.

**Supplementary Results and discussion**

**2.1 Bioaugmentation assays**

The bioaugmentation procedure began on day 132 (Fig. S5). Following this, *Ca.* Digestoria delfossei OTU_1 successfully established itself within the microbial community of the bioaugmented reactor. Notably, propionate consumption started within just a few days of starting the bioaugmentation, leading to a gradual pH increase. By day 146 (14 days after the procedure began), the reactor pH had returned to neutral, and the biogas produced contained over 50% CH₄. This restoration of anaerobic digestion demonstrates the effectiveness of bioaugmentation in mitigating reactor acidosis.

**2.2 Correlation network analysis**

The dataset generated in this study merged with the dataset from (Lemaigre et al., 2023) included 282 samples for a total of 8955 ASVs. Around 98.2% of these ASVs were from bacterial origin, while archaea represented only 1.8% of the studied community. Only 154 ASVs were kept after the first filtering (*i.e.* representing more than 0.1% of the total microbial community) for the next step (representing 1.7% of the initial number of ASVs). After the Spearman’s pairwise rank correlation calculation, a total of 2369 interactions having an adjusted *p*-value ≤ 0.001 and a R coefficient ≥ 0.5 (1709 positives (72.1%) and 660 negatives (27.9%)) were established between the previously selected 154 ASVs (100 were of bacterial origin (64.9%) and 54 of archaeal origin (35.1%), Fig S4A).

The Cloacimonadota OTU_1 was included in a direct neighbor correlation network (Fig. S4B) including 35 ASVs (26 from bacterial origin (74.3%) and 9 from archaeal origin (25.7%)) and 309 interactions ((238 positives (77.0%) and 71 negatives (23.0%)). Topological features, such as Degree, Betweenness, Closeness and Eigenvector centrality were used to identify potential specific keystone ASVs (Supplementary Dataset 1, Table S4). Interestingly, the first ASV highlighted as potential key partners of the *Ca.* Digestoria delfossei OTU_1 was from archaeal origin (A_ASV4) and taxonomically affiliated to *Methanothrix soehngenii*. A second one (A_ASV11) is taxonomically affiliated with the *Methanosarcina* genus.

The same approach was applied to the year-long monitoring dataset from (Calusinska et al., 2018) (Fig. S4C and D; details of the topological features are not included here). Using this independent dataset, a representative of the Methanothrix genus (archaeal OTU_3 from Calusinska *et al.*, 2018) and a member of the archaeal Bathyarchaeota phylum (archaeal OTU_15 from Calusinska *et al.*, 2018) again emerged as potential key partners of the Cloacimonadota_OTU_1.

**2.3 The presence of other, potentially novel syntrophic propionate oxidizing bacteria in the ABR1**

To ascertain the potential presence of other putative syntrophic propionate oxidising bacteria (SPOB) in the test reactor, we conducted analysis of the genomes of other bacteria within ABR1. We identified two metagenome assemble genomes (MAGs), including bin30 and bin45, both classified within the Bacteroidota, which encoded a potentially complete methyl malonyl-CoA (mmc) pathway, suggesting they may represent previously uncharacterized SPOB (Supplementary Dataset 3, Table S14; Fig. S8). However, most of the associated genes were scattered throughout the genome and not organized into a typical *mmc* cluster, as previously described for common SPOB (Westerholm et al., 2022). Additionally, a few other MAGs encoded components of the Cloacimonadota-specific *mmc* pathway (Fig. S8). Given the presence of these microbes, we sought to determine whether the novel *Ca.* Digestoria delfossei OTU_1 would still play a role in propionate consumption and whether it might employ a yet-undescribed metabolic pathway (see below). To address this, we briefly analyzed the metatranscriptomic abundance of known mmc genes across the different MAGs, including the novel *Ca.* Digestoria delfossei OTU_1. We observed that while the *mmc* genes were expressed by various MAGs during different experimental phases, their overall expression levels were very low (Supplementary Dataset 3, Table S14). This may be attributed to the disproportionately high expression of a few archaeal genes involved in methane production, which accounted for a significant proportion of the mapped reads in our metatranscriptomic dataset (Fig. S7). Consequently, much deeper sequencing would be required to adequately capture the expression patterns of genes with lower metatranscriptomic abundance. As a result, the metatranscriptomic analysis provided only limited insights, primarily confirming that all presumed SPOB expressed their *mmc* genes at some point during the experiment. A more detailed investigation of these other SPOB was beyond the scope of this study.

**2.4 Database of Cloacimonadota genomes and AD type-specific separation of Cloacimonadota clades**

With the exception of a few MAGs, the distribution of Cloacimonadota seems to be limited to the sea sediments and terrestrial methanogenic habitats only, *e.g.* anaerobic digestion reactors, sedimentary rocks and the termite gut (Supplementary Dataset 2, Table S7). Clades I (represented by 42 MAGs), III (15 MAGs) and IV (22 MAGs) contained species representative of AD and WWTP environments, at the same time regrouping the highest number of reconstructed MAGs. Clade IIc contained 34 MAGs of marine origin, including deep sea sediments and hydrothermal vents. It regrouped the highest diversity of candidate species (17 genome clusters - GCs) that were mainly represented by single MAGs (12 GCs). Two smaller clades contained MAGs originating from the termite gut studies (clade IIb; 3 MAGs) and ground water and sedimentary rocks (clade IIa; 5 MAGs), both regrouping the lowest number of MAGs, entirely representing separate candidate species (i.e. genome clusters). Genome size distribution (between 1.2 to 4 Mb), number of encoded proteins (1.2 to 3.4 k) and the GC content (32 to 56%) highlighted a significant genetic variation within Cloacimonadota (Fig. S9). However, no clear clade-specific trends were revealed.

Clades I and III were both representative of different anaerobic digestion systems. However, further 16S rRNA gene comparison (whenever the 16S rRNA gene was present in a MAG) with the previous study analyzing the abundance and stability of microbial communities in full scale energy units (Calusinska et al., 2018), showed that species from clade I were commonly abundant in different types of AD units. While Cloacimonadota from clade III dominated mainly ADs operating at WWTPs. This apparent separation is in accordance with the source of metagenomes that were at the origin of the different MAGs (Supplementary Dataset 2, Table S7).

**2.5 Protein clusters and hypothetical proteins in Cloacimonadota genomes**

Over half of the proteins could not be functionally annotated through whole-sequence searches against existing databases (Supplementary Dataset 7, Table S8), limiting our ability to perform an unbiased comparison of Cloacimonadota metabolic capacities. To address this, we grouped homologous proteins by clustering over 200,000 proteins extracted from the analyzed Cloacimonadota genomes into protein clusters (PCs). This clustering yielded 21,260 PCs, including 12,043 singletons (representing 5.7% of all proteins), which are isolated proteins with no homologs in distinct Cloacimonadota genomes. Using MAGs as nodes (colored by their origin) and correlations as edges, we assessed the discriminative power of PCs by generating a network (Fig. S10). MAGs from different tree clades (Fig. 2) clustered according to habitat, separating into three main groups: the termite gut, anaerobic digesters, and marine sediments (Fig. S10A). Marine MAGs appeared more dispersed in the network compared to those from the other two habitats, indicating significantly higher genetic diversity within this clade. This observation aligns with the rarefaction curve, which revealed a much greater diversity of Pfam domains in marine Cloacimonadota (Fig. S10B), potentially reflecting a broader metabolic repertoire. Interestingly, accumulation curves for unique Pfam domains within each clade showed near saturation for AD-associated clades (I, III, and IV), plateauing at around 2,500 unique domains, with only a minimal increase at the curve's end. This suggests that further genome sequencing is unlikely to uncover many additional domains for these clades. In contrast, the marine cluster (IIC) exhibited a steep curve that did not plateau even after analysing 34 genomes, highlighting the vast unexplored metabolic potential of marine Cloacimonadota.

To further investigate functional diversity, PCs were assigned to KEGG Orthologies (KOs) to evaluate subfamily variation among functionally assigned PCs. Overall, no correlation was observed between the diversity (number of PCs) associated with a KO category and the total number of genes assigned to that category. This suggests that while certain functions are evolutionarily conserved within the Cloacimonadota phylum, others exhibit considerable variation (data not shown). Among the top 100 most abundant KOs, 20 were represented by single PCs, nearly half by two PCs, and over 80% by just five PCs (Supplementary Dataset 2, Table S10). We hypothesize that functions encoded by proteins assigned to the most abundant KOs but represented by single PCs are likely of fundamental importance to the phylum's activity. These functions, being widespread yet highly conserved (i.e., unchanged at the sequence level), likely play a critical role in Cloacimonadota metabolism. For example, three of the most abundant KOs represented by single PCs include phosphate butyryltransferase (likely propionyltransferase; see main text for details), methylmalonyl-CoA decarboxylase subunit alpha, and the type IV pilus assembly protein PilB. The first two are components of a propionate oxidation pathway, while the presence of PilB suggests a potential role in direct electron transfer (DIET), a characteristic also observed in some SPOB (Westerholm et al., 2022). Additional genes involved in the methylmalonyl-CoA pathway were among the most abundant and conserved PCs, further supporting the hypothesis that propionate oxidation is a phylum-wide capability of Cloacimonadota.

In contrast, conserved housekeeping genes, such as *rpoD* (encoding RNA polymerase primary sigma factor), exhibited less conservation within Cloacimonadota, being represented by seven PCs. KOs with the highest PC diversity (over 20 PCs) included two putative transposases (K07497 and K07491, enriched in AD and marine Cloacimonadota MAGs, respectively), two uncharacterized protein groups (K07126 and K07076, again enriched in AD and marine MAGs, respectively), eukaryotic-like serine/threonine-protein kinase K12132, and the type I restriction enzyme S subunit K01154. Although the latter was represented by only 55 genes, it exhibited high diversity (26 PCs) and was enriched in AD MAGs. These findings highlight significant functional diversity among Cloacimonadota, particularly between different habitat types.

A total of 3,664 PCs, representing 49.2% of proteins, were assigned KOs functionalities, leaving a substantial portion of proteins classified as hypothetical. Given the well-established fact that protein structures are often more conserved than their sequences, we further analyzed Pfam domains. Based on the premise that proteins with similar structures are likely to share similar biological functions, we aimed to gain further insights into the functional potential of these uncharacterized proteins. To evaluate whether hypothetical proteins might harbor novel metabolic functions, we compared the distribution of Pfam domains between functionally assigned and unassigned proteins. The analysis revealed a weak correlation (Pearson’s correlation coefficient, r = 0.36), suggesting that the metabolic potential encoded by hypothetical proteins may include entirely new functions. This finding underscores the possibility of uncovering novel biochemical pathways and metabolic activities within the unexplored protein pool.

**2.6 Carbohydrate active enzymes in Cloacimonadota genomes**

Previously, Cloacimonadota from engineered environments were found to encode a larger repertoire of carbohydrate active enzymes (CAZymes) compared to those in natural systems (Johnson & Hug, 2022). However, these primarily include multiple glycoside transferase coding genes (Supplementary Dataset 2, Table S11). In contrast, the diversity and number of hydrolysis CAZyme gene copies are much lower in Cloacimonadota compared to, for example, Bacteroidota or Bacillota (former Firmicutes) (Campanaro et al., 2020), suggesting their contribution to carbohydrate degradation in AD systems is rather limited. In clade I AD Cloacimonadota, multiple CAZymes are organized into two gene complexes which respectively target starch (α-glucans) and mannose (including acetyl-D-hexosamine) residue-containing carbohydrates. In the case of clade IIA, the termite clade IIB, and the other AD clades III and IV, only the former CAZyme complex was present. For comparison, no specific CAZyme gene complexes were identified in most of the marine Cloacimonetes. However, the reconstructed genomes were more fragmented, which might have impacted the search results.

**2.7 Enrichment and cultivation of Claocimonadota from full-scale anaerobic digestion reactors**

**Enrichment and cultivation trials**

Based on 16S rRNA analysis, one enrichment (RT_BT_as_ph7; Supplementary Dataset 5, Table S18) achieved a high dominance of a single OTU belonging to Cloacimonadota, comprising 56% of its relative abundance (Supplementary Dataset 5, Table S19). This enrichment was cultivated in PYGV medium (DSMZ 621 basal medium) supplemented with ampicillin (100 mg/L), streptomycin (100 mg/L), and D-glucose (1% w/v). Subsequent re-culture on PYGV medium supplemented with propionate (10% w/v) further stimulated the growth of the Cloacimonadota OTU (Supplementary Dataset 5, Table S20). This sample was subsequently used for metagenomic reconstruction (Supplementary Dataset 5, Table S21). Sequencing and analysis revealed that the enriched Cloacimonadota OTU represented a distinct genomic species (GS28 in Fig. 2), which we later named *Ca.* Cloacimonas fortuita. Additionally, the most abundant archaeal species in the enrichment was *Methanothrix* sp. (Fig. 5), supporting its putative role as a syntrophic partner of Cloacimonadota in AD reactors (Supplementary Dataset 1, Table S4).

Despite repeated transfers of the enriched culture to fresh medium, enriched Cloacimonadota cells were consistently lost (significant decrease in abundance), suggesting the absence of an essential factor present in the original sludge. To optimize conditions for isolation, 14 different media formulations and growth conditions were further tested, informed by genomic analyses of Cloacimonadota functional traits in AD environments. Additives such as acetate, propionate, pyruvate, potassium chromate, magnetite, and L-glutamate were included as potential growth promoters (Supplementary Dataset 5, Table S20). However, these efforts did not yield successful isolation of the target species (details not discussed here). The failure to isolate *Ca.* Cloacimonas fortuita may be attributed to stochastic fluctuations or the growth dependency on its archaeal symbiont *Methanothrix*, which appears crucial for its survival. Other enrichments cultivated in the tested media and conditions also contained OTUs belonging to Cloacimonadota (Supplementary Dataset 5, Table S19), but none reached the high abundance observed in the dominant OTU enrichment. These results highlight the intricate ecological and physiological requirements of Cloacimonadota and underscore the challenges associated with isolating and maintaining members of this phylum in laboratory conditions.

**Designation of two new *Cloacimonadota* species**

***Candidatus* Cloacimonas fortuita**

The genome of the enriched bacterium in the culture RT_BT_as_ph7 was analyzed and compared to the known *Ca.* Cloacimonas acidaminovorans (GCA_000146065.1). Genomes of the enriched species and the closely related species *Ca.* C. acidaminovorans have an ANI of 78.3%, which is below the species-level cut-off (95%) but above the genus-level threshold (>70%). Their AAI value of 79% further supports that they belong to the same genus, *Ca.* Cloacimonas. Therefore, we propose the newly recovered genome as the nomenclatural type for the novel species within the Cloacimonas genus, and name it as *Candidatus* Cloacimonas fortuita sp. nov. This species has been registered in SeqCode (<https://seqco.de/i:52907>).

*Candidatus* Cloacimonas fortuita (for.tu.i'ta, L. fem. adj. fortuita, accidental, fortuitous, by chance; referring to the accidental isolation of the strain).

***Candidatus* Digestoria delfossei**

The genome of the enriched bacterium in the ABR1 reactor was analyzed and compared to its closely related species, previously described *Ca.* Syntrophosphaera thermopropionivorans (GCA_004353895.1). ANI and AAI comparative analyses revealed values of 66.5% ANI and 66.4% AAI, respectively. These results surpassed the species-level threshold of 95% ANI while placing the bacterium beyond the genus-level range of AAI (below 70%). Additionally, both genomes have largely different GC content, which is usually conserved within the same genus, reinforcing the likelihood that genome of the enriched bacterium represents a new genus. However, due to the lack of precise AAI cut-offs for *Ca.* Cloacimonadota, further studies, including full-length 16S rRNA comparisons, should complement these findings. Together, these findings indicated that the enriched Cloacimonadota species likely represent a new species within a genus distinct from *Ca.* S. thermopropionivorans. Based on genomic analysis, we propose it serves as the nomenclatural type for the novel species and genus *Candidatus* Digestoria delfossei gen. nov. sp. nov. This species has been registered in SeqCode (<https://seqco.de/i:52909>). All associated higher taxa are left unnamed at this time, pending further resolution of the taxonomy of the phylum Cloacimonadota.

*Candidatus* Digestoria gen. nov. (Di.ges.to´ri.a. L. fem. n. digestio, digestion; N.L. fem. n. Digestoria, a genus name referring to its role in anaerobic digestion).

*Candidatus* Digestoria delfossei sp. nov. (del.fos’se.i. N.L. gen. n. delfossei, named in honor of our former group leader, Dr. Philippe Delfosse).

**Supplementary Figures**

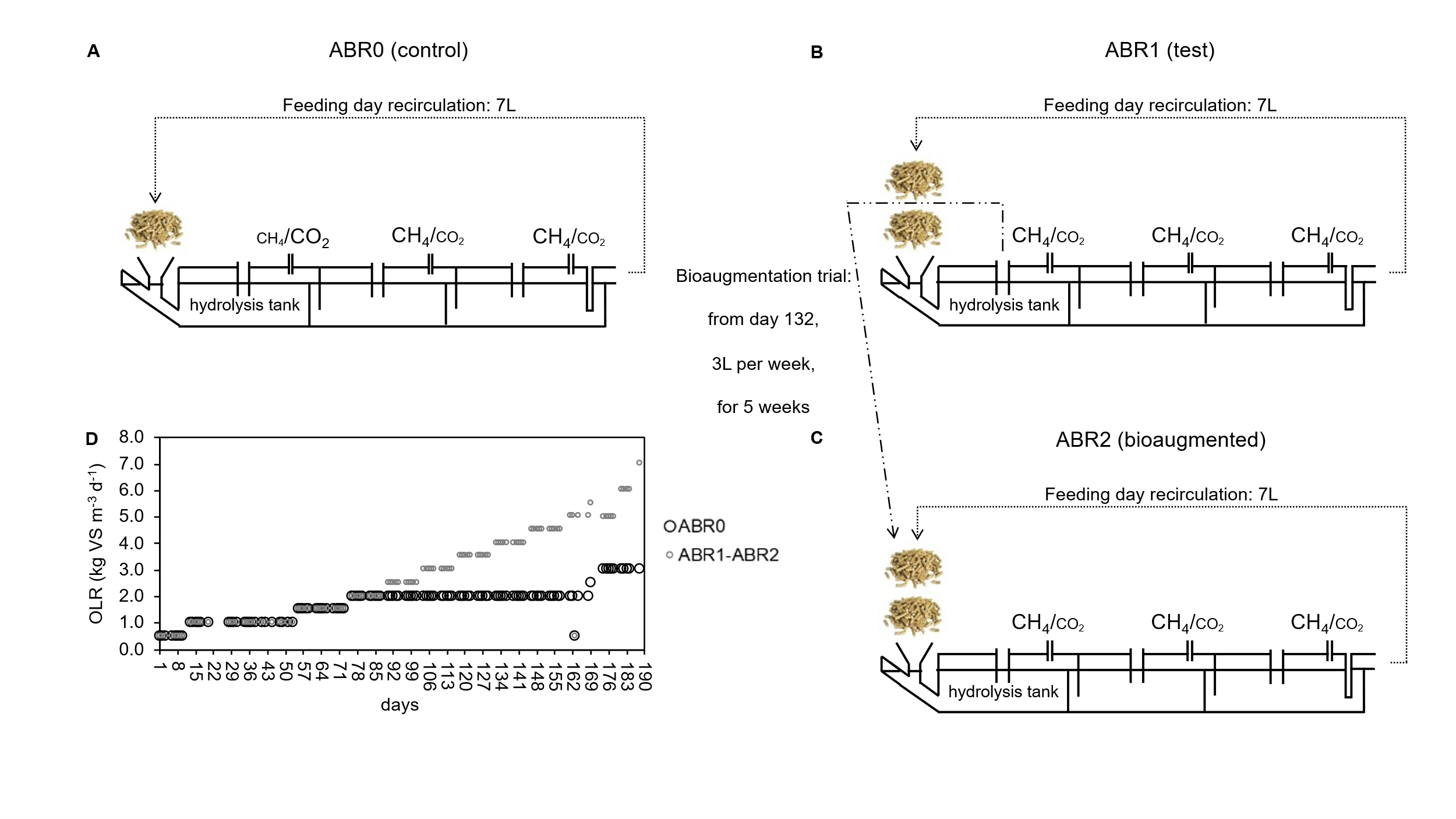

**Figure S1.** A detailed feeding and sludge recirculation plan implemented for the anaerobic baffled reactors (ABRs), encompassing the control reactor (ABR0; A), the test reactor (ABR1; B), and the bio-augmented reactor (ABR2; C).

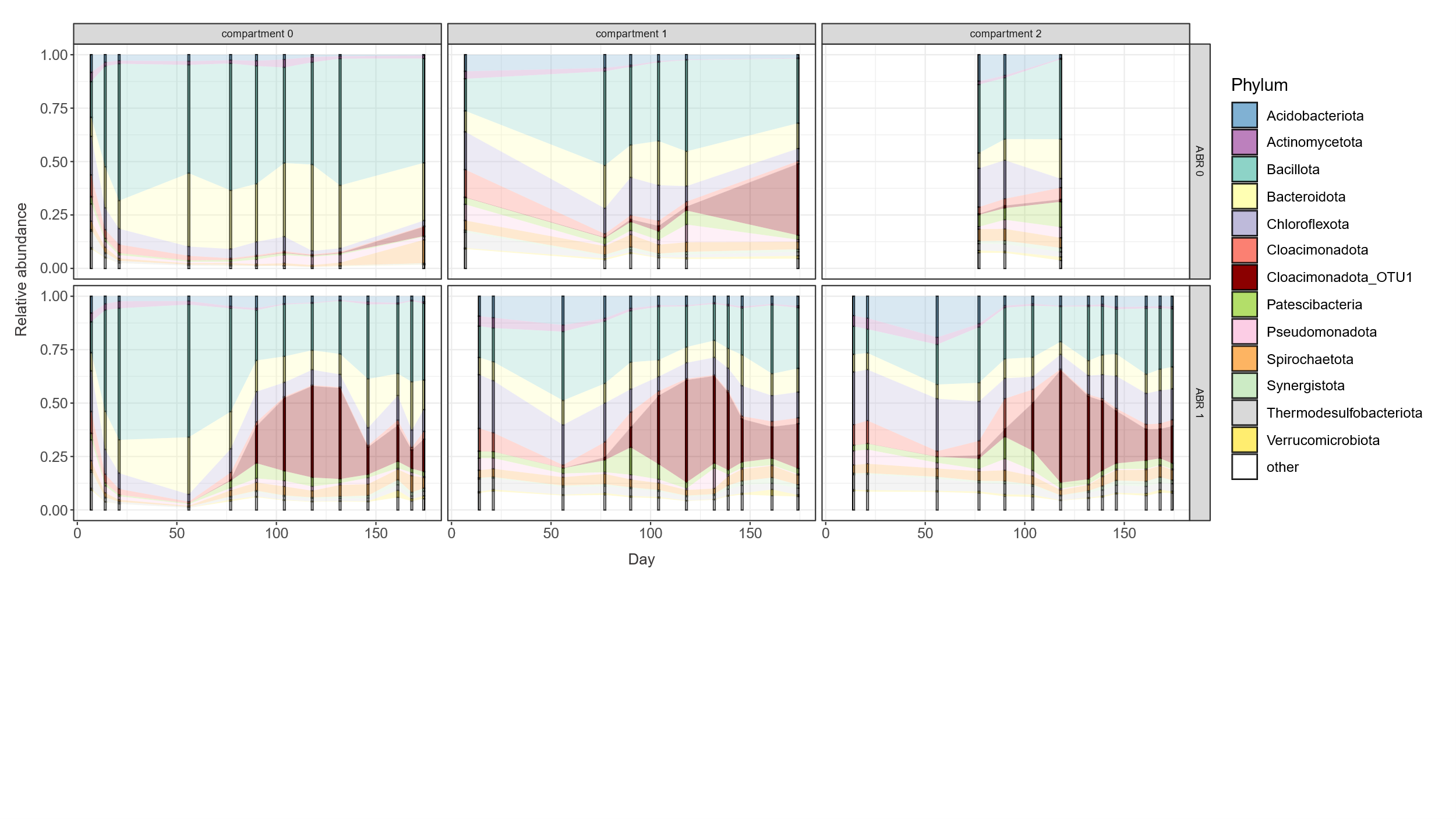

**Figure S2.** Phylum-level bacterial community composition in the control reactor (ABR0) and the test reactor (ABR1) across the different reactor compartments, as determined by 16S rRNA gene amplicon sequencing (Table S2).

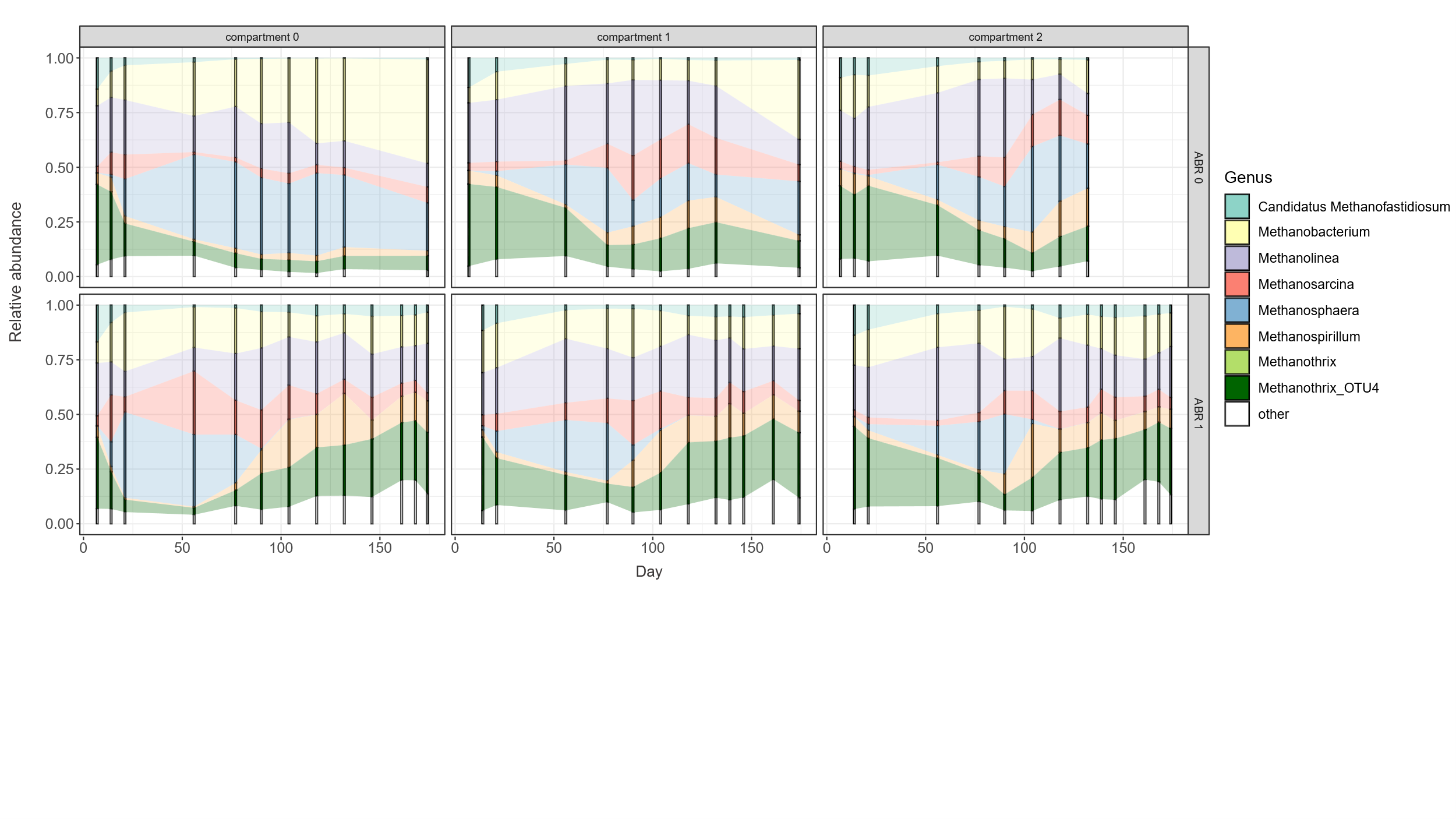

**Figure S3.** Genus-level archaeal community composition in the control reactor (ABR0) and the test reactor (ABR1) across the different reactor compartments, as determined by 16S rRNA gene amplicon sequencing (Table S3).

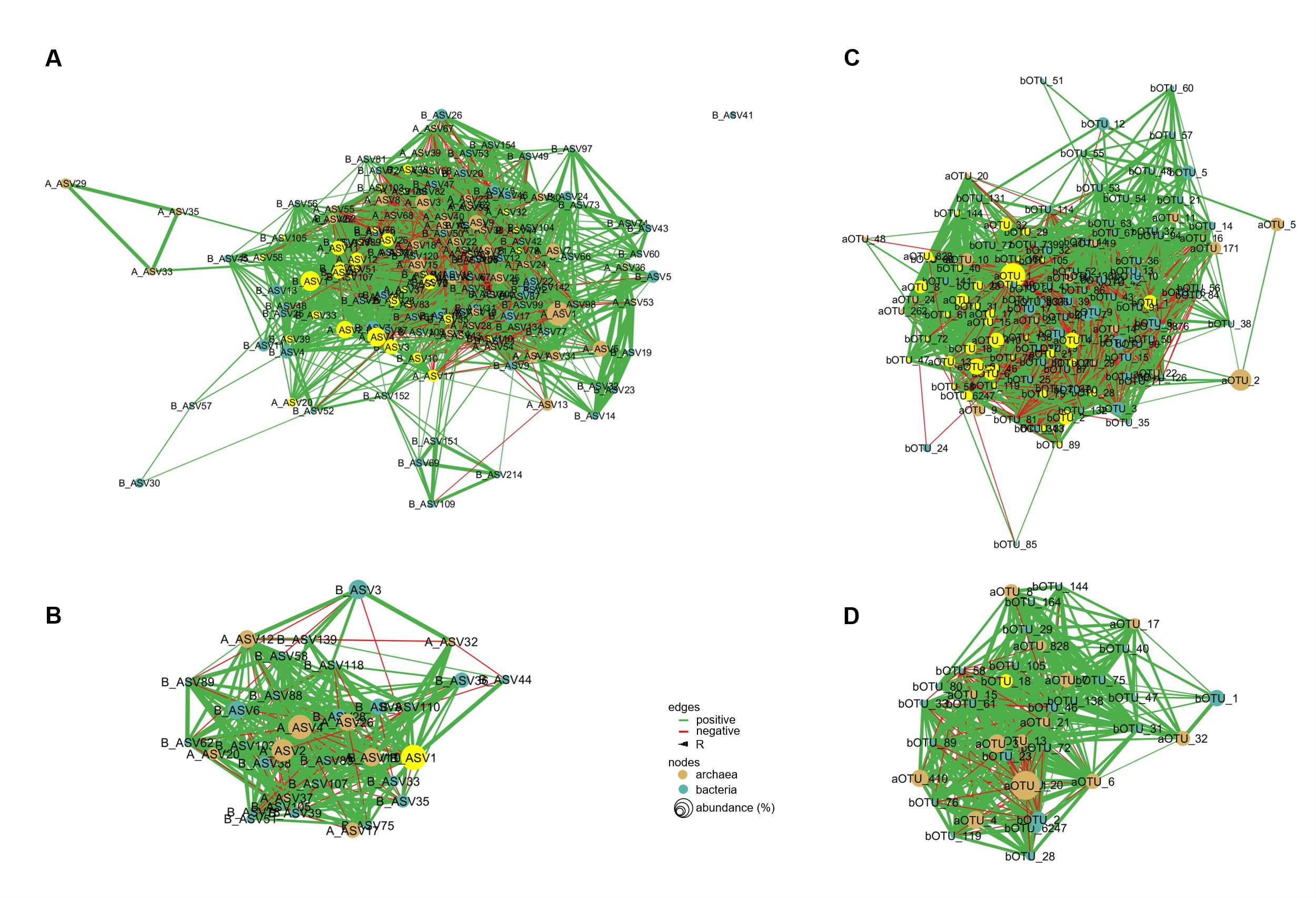

**Figure S4.** Correlation network analysis (CNA). General CNA (A) and direct 'Cloacimonadota OTU_1' (B; assigned as B_ASV1 on this figure) neighbour CNA, constructed for the dataset from this study merged with the dataset from (Lemaigre *et al.*, 2018; at the ASVs level). **C**. and **D**. General CAN (C) and direct 'Cloacimonadota OTU_1' (D; assigned as B_ASV1 on this figure) neighbour CNA, constructed for the dataset from the year-long monitoring study by (Calusinska *et al.*, 2018; at the OTU level). Only ASVs and OTUs representing more than 0.1% of the whole bacterial and archaeal community were used for the Spearman’s rank correlation calculation. Only highly significant correlations, *i.e.* with adjusted *p*-value ≤ 0.001 and R coefficient ≥ 0.5, are represented. Nodes in blue represent bacterial origins, while nodes in brown correspond to archaeal origins. Nodes in yellow in panels A and C represent the direct neighbors of 'Cloacimonadota OTU_1'. Nodes in yellow in panels B and D represent Cloacimonadota_OTU1, identified as B_ASV1 in panel B and bOTU_18 in panel D. Green edges and red edges, respectively, indicate positive and negative correlations. Edge thickness corresponds to the R coefficient value of the correlation, while node size reflects the relative abundance of ASVs or OTUs within the bacterial and archaeal communities of the respective dataset.

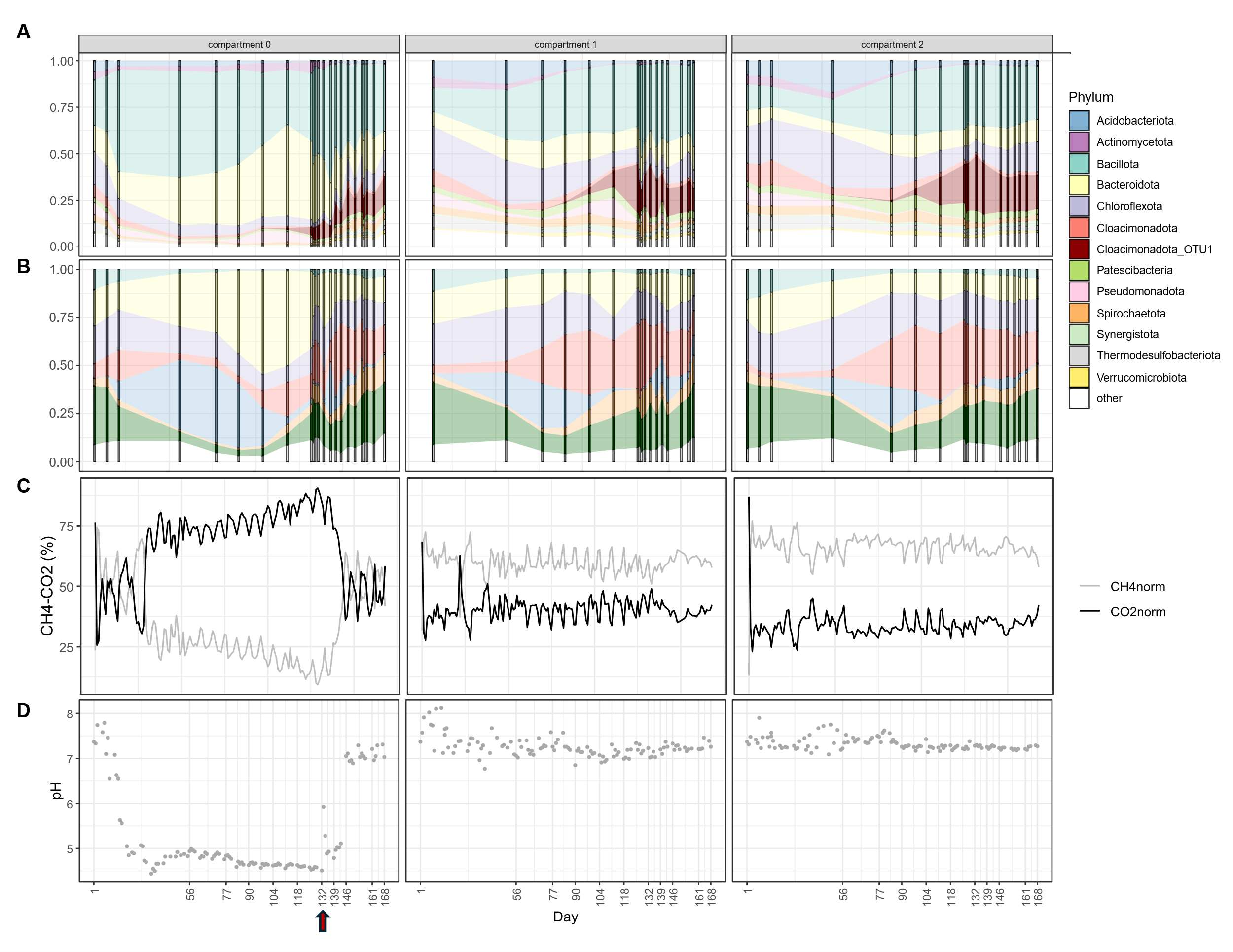

**Figure S5.** Bio-augmentation trials of the acidified anaerobic baffled reactor (ABR2) using Cloacimonadota OTU_1-enriched sludge. Phylum-level bacterial (A) and genus-level archaeal (B) community composition in ABR2 across the different compartments; percentages of CH₄ and CO₂ in the biogas produced by ABR2 (B); pH values and propionate concentrations in ABR2 (C). The black arrow marks day 132, indicating the start of bio-augmentation with sludge enriched in *Ca.* Cloacimonadota OTU_1.

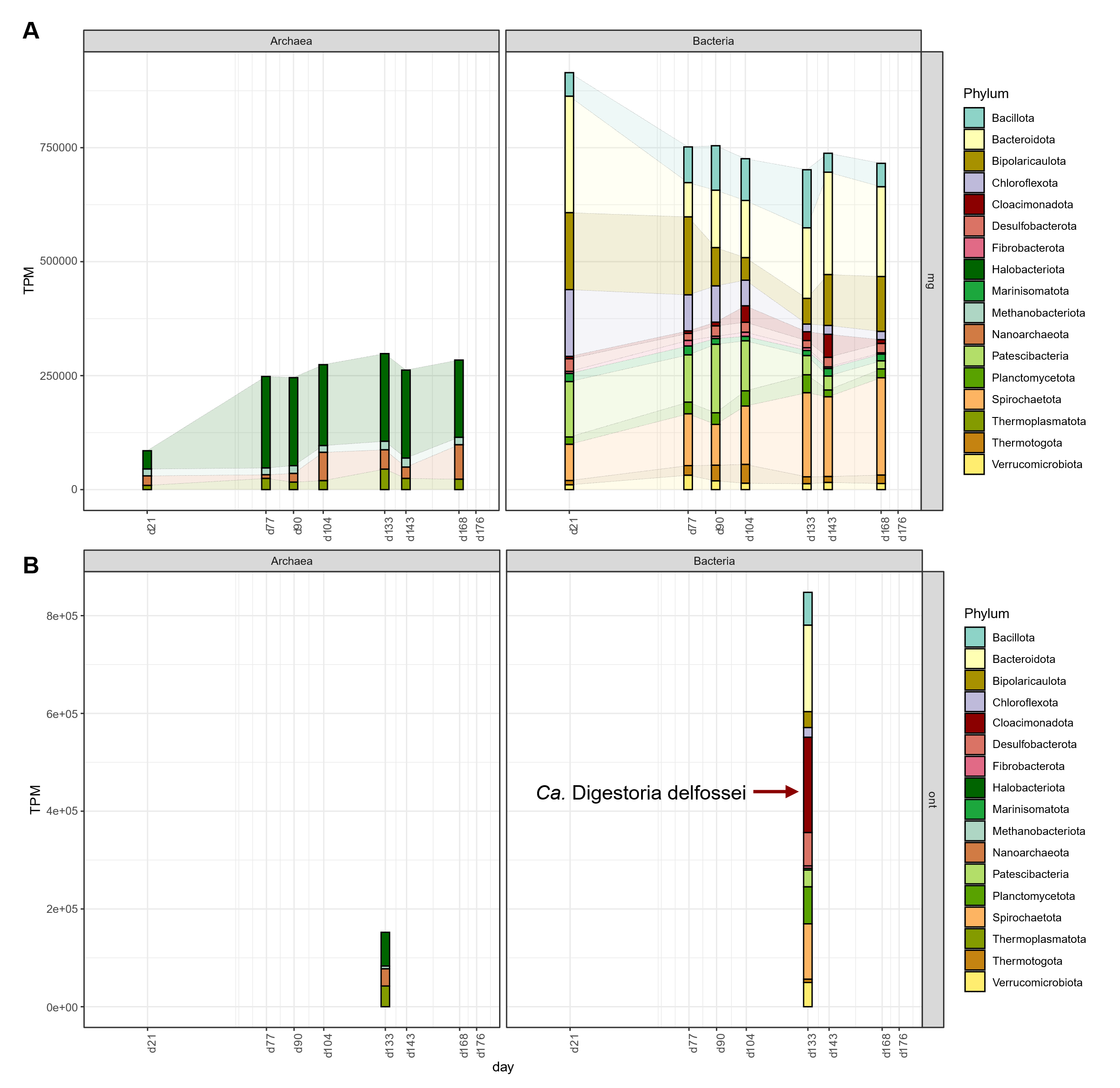

**Figure S6.** Phylum-level metagenomic abundance over time of metagenome-assembled genomes (MAGs) reconstructed in this study, based on short (A) and long (B) read sequencing. The genome of *Ca.* Digestoria delfossei, corresponding to OTU_1 in Fig. 1, is highlighted.

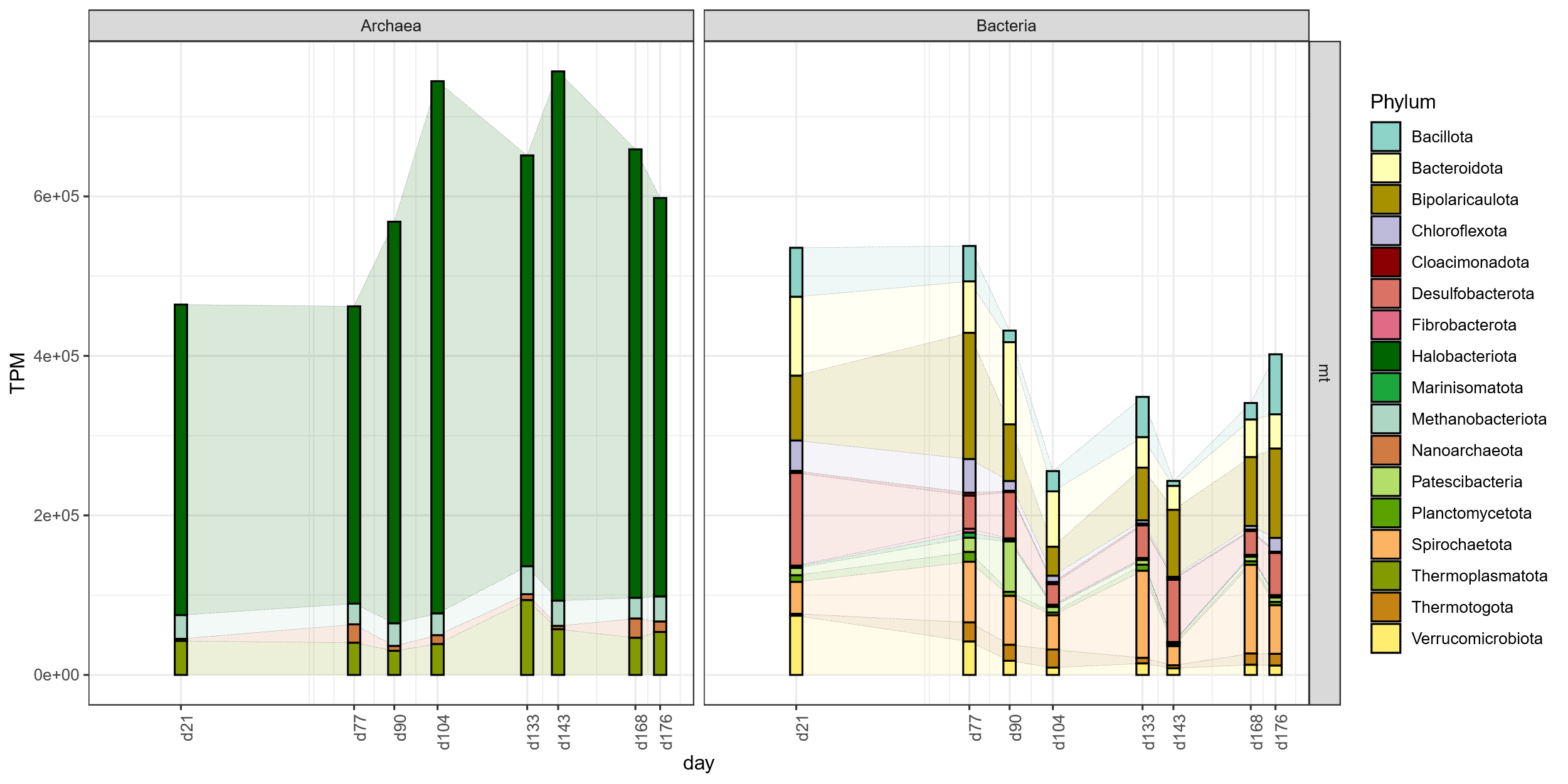

**Figure S7.** Phylum-level metatranscriptomic abundance over time of metagenome-assembled genomes (MAGs) reconstructed in this study (A). High metatranscriptomics abundance of archaeal MAGs results from very high gene transcript abundance of genes involved in the methanogenesis pathway (B).

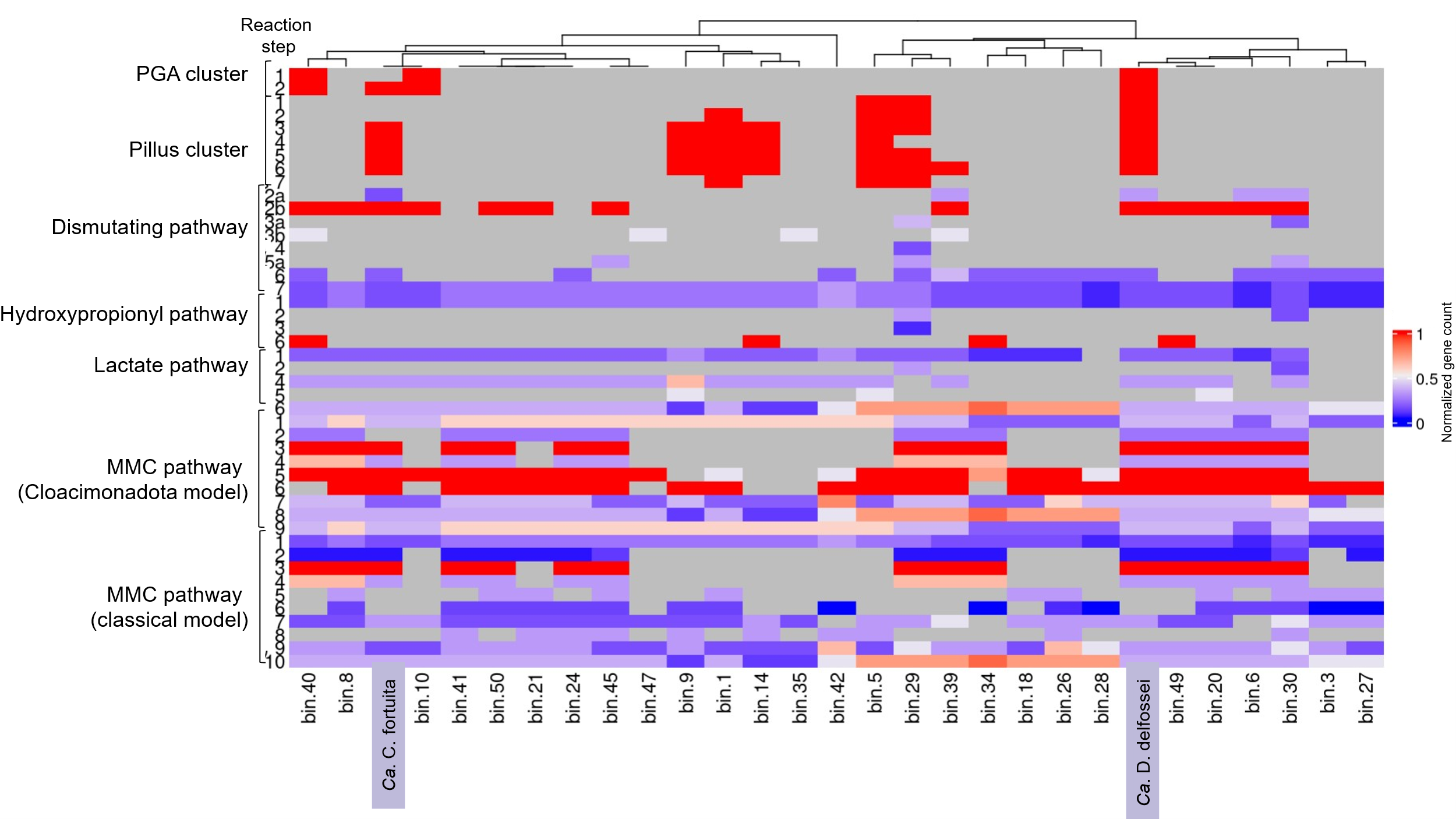

**Figure S8.** Potential syntrophic propionate-oxidizing bacteria (SPOB) based on pathway completion in the metagenome-assembled genomes generated in this study (Table S12).

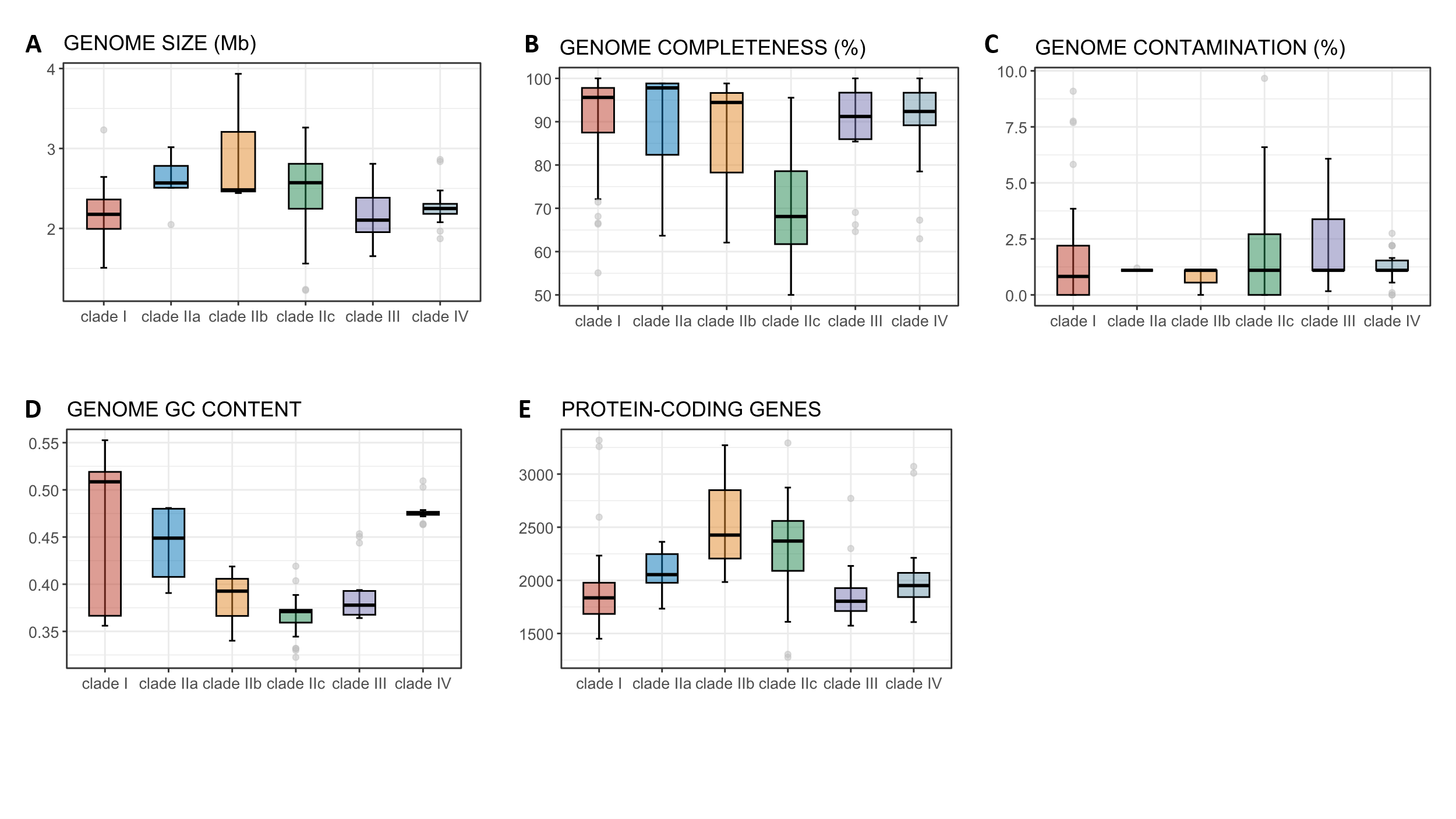

**Figure S9.** Characteristics of Cloacimonadota genomes used in the final database at the tree clade level (Fig. 2), including average genome completeness (A), average genome contamination (B), average GC content per genome cluster (C), average genome size per genome cluster (D; extrapolated for less complete genomes), and number of protein-coding genes (E; extrapolated for less complete genomes).

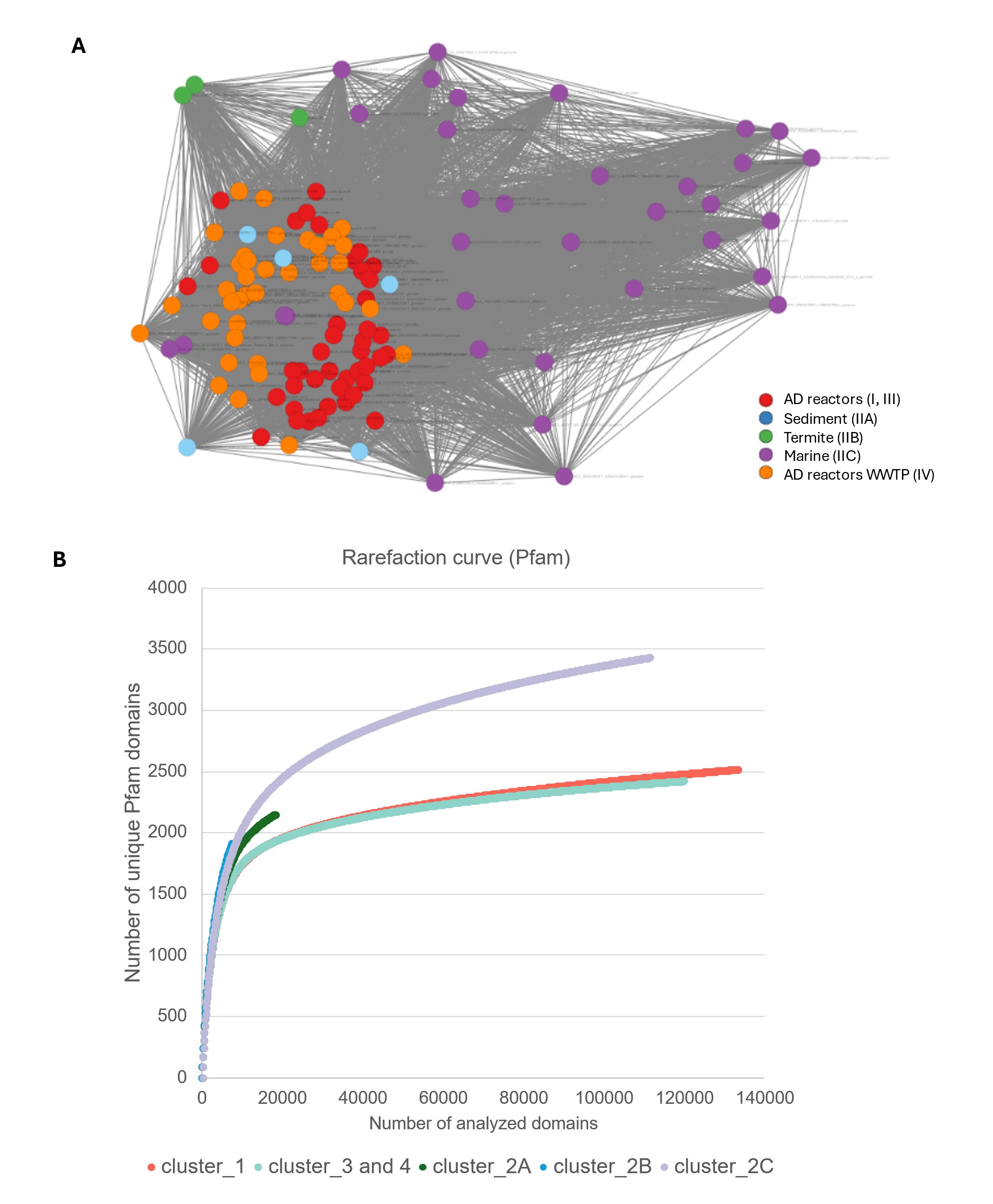

**Figure S10.** Protein cluster (PC) network (A) and Rarefaction curves of unique Pfam domains versus the number of analyzed Pfam domains (B) generated for different Cloacimonadota genomes, classified by tree clade (Fig. 2). These curves provide insights into the diversity and functional richness of Pfam domains within each cluster, highlighting variations in domain representation across genomes.

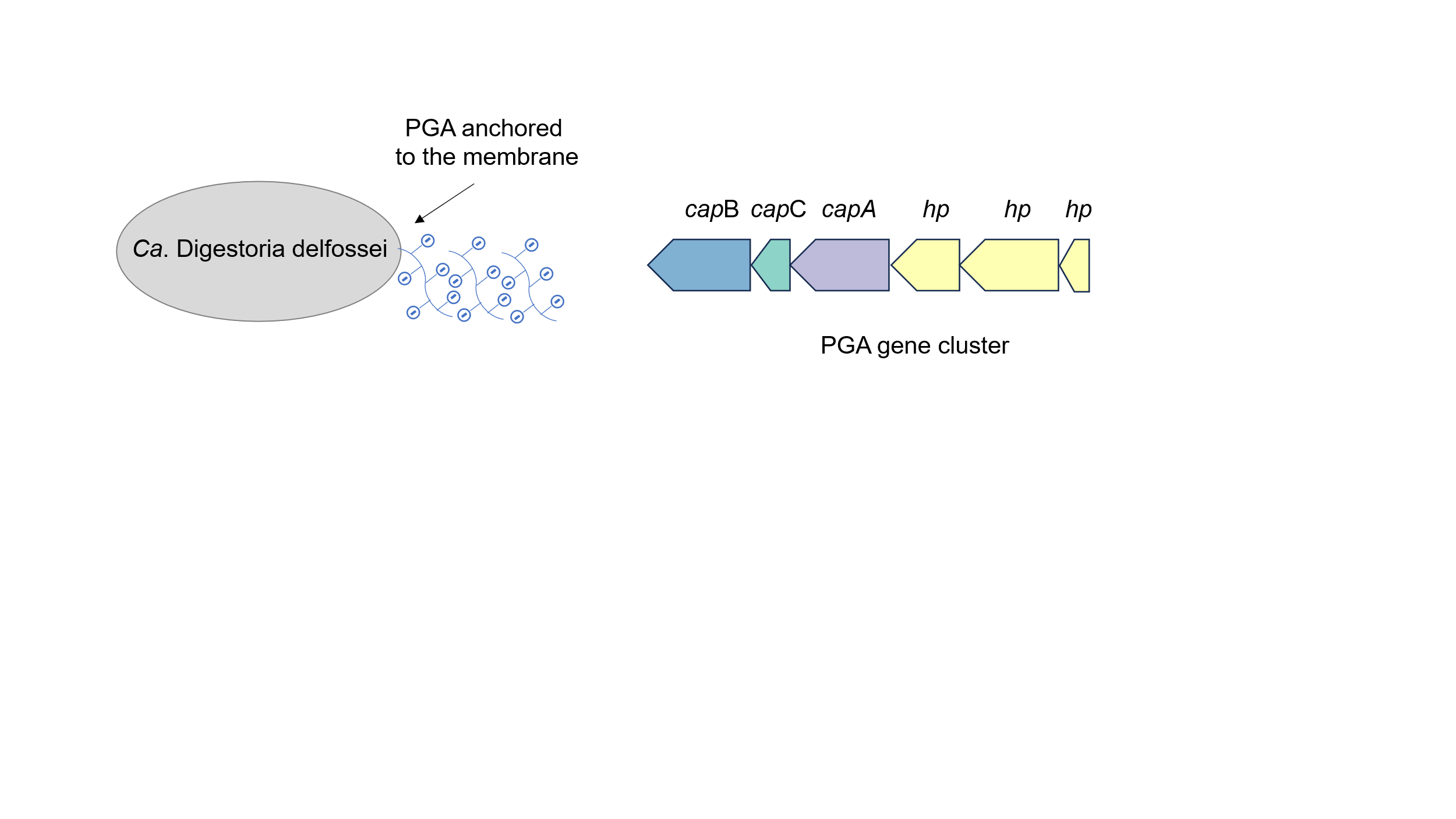

**Figure S11.** Gene organization in a poly-ɣ-glutamate (PGA) cluster within the *Ca*. Digestoria delfossei genome. PGA – poly-ɣ-glutamate; hp – hypothetical

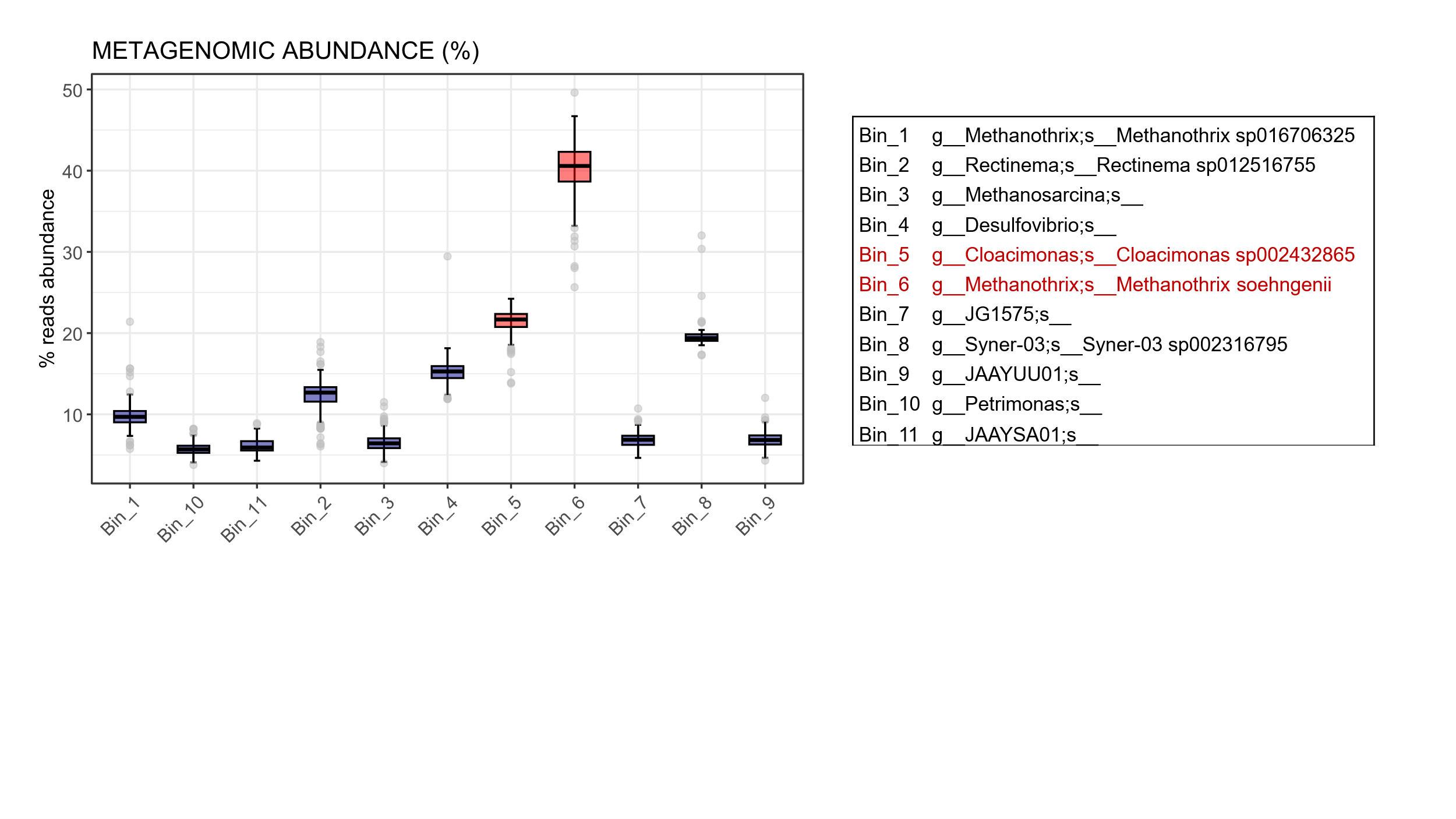

**Figure S12.** The metagenomic abundance (% of reads) of species (bins) identified in the Cloacimonadota-enriched culture grown on propionate (10 g/L) was analyzed, along with their taxonomic classification using the GTDB-Tk database.

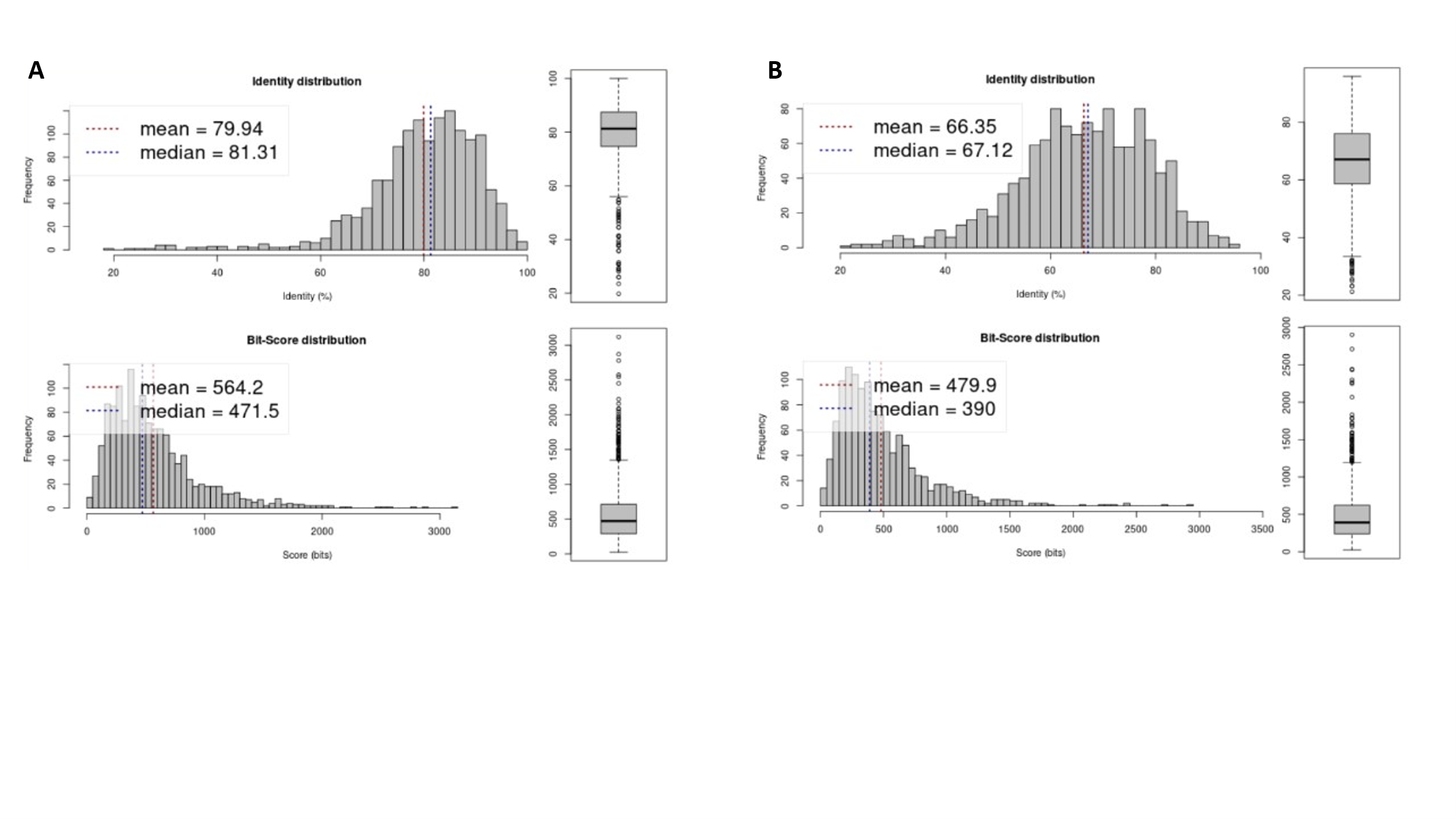

**Figure S13.** Amino acid identity (AAI) scores for shared proteins between *Ca*. Cloacimonas fortuita and *Ca*. Cloacimonas acidaminovorans (A) and *Ca.* Digestoria delfossei and *Ca.* Syntrophosphaera thermopropionivorans (B). The top panel shows the percentage identity distribution, indicating the level of amino acid similarity between the compared genomes. The bottom panel displays the bit-score distribution, reflecting the statistical significance of the alignments.

**Description of Supplementary Datasets and Tables**

**Supplementary Dataset 1: Anaerobic baffled reactors (ABR), experimental design and 16S rRNA gene amplicon sequencing.**

**Table S1.** Lab-scale anaerobic baffled reactors (ABRs) and measured variables.

**Table S2.** Count table of the 16S rRNA gene amplicon sequencing for bacteria. Reads were normalized to 8,000 per sample. Samples marked with * are the samples presented in Fig. 1.

**Table S3.** Count table of the 16S rRNA gene amplicon sequencing for archaea. Reads were normalized to 5,000 per sample. Samples marked with * are the samples presented in Fig. S3.

**Table S4**. Topological features for ASVs involved in the direct "cloacimonadota OTU_1" (Ca. Digestoria delfossei) neighbor correlation network for the dataset of this study merged with the dataset from Lemaigre et al., 2018.

**Supplementary Dataset 2: Genome reconstruction of microbes present in Anaerobic Baffled Reactor 1 (ABR1) and the Cloacimonadota genome database.**

**Table S5**. Characteristics of metagenome-assembled genomes (MAGs) reconstructed in this study from Illumina short-read metagenomics (before genome refinement for bin36 representing the *Ca*. Digestoria delfossei).

**Table S6.** Details of the two reconstructed Cloacimonadota genomes from species enriched in this study.

**Table S7.** Complete list of 121 Cloacimonadota metagenome assembled genomes (MAGs) used in the final Cloacimonadota database, including MAGs generated in this study and from other sources (downloaded in May 2020).

**Table S8.** Characteristics of KEGG orthologues (KO) assignments in genomes included in the final Cloacimonadota database. The numbers indicate the presence of specific KOs in genomes belonging to the Cloacimonadota clade (as shown in Fig. 2). Linear Discriminant Analysis Effect Size (LEfSe) was employed to identify features (i.e., KOs) that best explain differences between clades. The analysis was performed using the Galaxy server (<https://huttenhower.sph.harvard.edu/lefse/>).

**Table S9.** Core metabolism of Cloacimonadota, inferred from KOs conserved across all Cloacimonadota clades (referenced in Fig. 2).

**Table S10.** Protein clusters (PCs) generated by MMSeq2 (Mirdita et al., 2019) and their functional assignment to KEGG orthologues (KOs) for the Cloacimonadota genomes included in the final database (Table S7).

**Table S11.** Summary of the carbohydrate-active enzymes (CAZy) domains identified by dbCAN2 (Zhang et al., 2018) for the Cloacimonadota genomes included in the final database (Table S7) and summarized at the tree clade level (Fig. 2).

**Supplementary Dataset 3: Syntrophic propionate oxidation (SPO) pathway and supporting results**

**Table S12.** KEGG orthologues (KOs) assignments to the various SPO pathways analyzed in this study. The hydroxypropionyl and lactate SPO pathways are adapted from Paton et al., 2020. Details on the alternative mmc pathway in Cloacimonadota is given in Table S9.

**Table S13.** Reaction steps and enzymes involved in the proposed alternative mmc pathway in Cloacimonadota.

**Table S14.** Genes associated with KOs involved in the steps of syntrophic propionate oxidation pathways (Tables S1 and S13) and assigned to the MAGs generated in this study (Table S5). Gene expression levels are presented as TPMs, with raw read counts also included.

**Table S15.** Proteins identified through peptides from the metaproteomic study, involved in the mmc pathway (Cloacimonadota alternative) or associated with SPO, are highlighted. The search was performed against the complete genome of the newly reconstructed *Ca.* Digestoria delfossei.

**Supplementary Dataset 4: Distinct and overlapping metabolic capacities of Cloacimonadota compared to other phyla in anaerobic digestion (AD) reactor**

**Table S16.** Representation of KEGG orthologues (KOs) in the genomes of Cloacimonadota (Table S7; limited to genomes from AD microbes) and other microbes within the AD microbiome (Campanaro et al., 2020). Only genomes with at least 70% genome completeness are included.

**Table S17.** Representation of Pfam domain in the genomes of Cloacimonadota (Table S7; limited to genomes from AD microbes) and other microbes within the AD microbiome (Campanaro et al., 2020). Only genomes with at least 70% genome completeness are included.

**Supplementary Dataset 5: Enrichment and cultivation trials of Cloacimonadota.**

**Table S18.** Summary of the conditions used to enrich *Ca.* Cloacimonadota from anaerobic digestion sludge (first stage). In bold, the culture with significant enrichment of *Ca.* Cloacimonas fortuita is shown.

**Table S19.** Screening of enriched cultures for the presence of novel Cloacimonadota species, with results expressed as the percentage of read abundance based on the 16S rRNA gene amplicon sequencing.

**Table S20.** Summary of the conditions used to enrich Ca. Cloacimonas fortuita from the RT_BT_as_ph7 culture initially enriched during the first stage (Table S19).

**Table S21.** List of species identified in the Cloacimonetes-enriched culture (*Ca.* Cloacimonas fortuita) grown RT_BT_as_ph7 on propionate (Clo3, Table S20) along with their genome characteristics. Genomes were reconstructed from short Illumina reads. *Ca.* Cloacimonas fortuitus bin is highlighted in bold.
